# Elucidating polymorphs of crystal structures with intensity-based hierarchical clustering analysis on multiple diffraction datasets

**DOI:** 10.1101/2022.09.13.507775

**Authors:** Hiroaki Matsuura, Naoki Sakai, Sachiko Toma-Fukai, Norifumi Muraki, Koki Hayama, Hironari Kamikubo, Shigetoshi Aono, Yoshiaki Kawano, Masaki Yamamoto, Kunio Hirata

## Abstract

In macromolecular structure determination using X-ray diffraction from multiple crystals, the presence of different structures (structural polymorphs) necessitates the classification of diffraction data for appropriate structural analysis. Hierarchical clustering analysis (HCA) is a promising technique that has so far been used to extract isomorphous data, mainly for single structure determination. Although in principle the use of HCA can be extended to detect polymorphs, the absence of a reference for defining a threshold used for grouping the isomorphous datasets (‘isomorphic threshold’) poses a challenge. Here, we have applied unit cell-based and intensity-based HCAs to the datasets of apo-trypsin and inhibitor-bound trypsin that were mixed post-data acquisition to investigate how effective HCA is in classifying polymorphous datasets. Single-step intensity-based HCA successfully classified polymorphs with a certain ‘isomorphic threshold’. In datasets of several samples containing an unknown degree of structural heterogeneity, polymorphs could be identified by intensity-based HCA using the suggested ‘isomorphic threshold’. Polymorphs were also detected in single crystals using the data collected by the continuous helical scheme. These findings are expected to facilitate the determination of multiple structural snapshots by exploiting automated data collection and analysis.

**Synopsis:** Single-step intensity-based hierarchical clustering is demonstrated to allow the detection of structural polymorphs in the diffraction datasets obtained from multiple crystals. By splitting the datasets collected by continuous helical scheme into several chunks, both inter and intra-crystal polymorphs can be successfully analyzed.

## 1. Introduction

Automation and acceleration of data collection at the macromolecular crystallography (MX) beamlines of synchrotron radiation facilities are yielding massive amounts of X-ray diffraction data. Such highly efficient, high yield data acquisition is now becoming a major trend in MX, namely, high data-rate MX (HDRMX) (Bernstein *et al*., 2020). HDRMX has been realized by the availability of highly brilliant X-ray beams (Ursby *et al*., 2020; Sanchez-Weatherby *et al*., 2019; Hirata *et al*., 2013), fast readout of detectors (Casanas *et al*., 2016), rapid sample exchange robots (Murakami *et al*., 2020; Papp *et al*., 2017; Martiel *et al*., 2020; Nurizzo *et al*., 2016), and the application of automated measurement schemes (Zander *et al*., 2015; Hirata *et al*., 2019; Basu *et al*., 2019; Bowler *et al*., 2016). Furthermore, data reduction and structure analysis of the obtained datasets has also been automated in various pipelines (Yamashita *et al*., 2018; Winter & McAuley, 2011; Wojdyr *et al*., 2013; Winter, 2009; Monaco *et al*., 2013; Vonrhein *et al*., 2011; Incardona *et al*., 2009; Wang *et al*., 2022). Finally, data management systems, including user interfaces for viewing these data, have been developed so that the experimenters can seamlessly manage and easily interpret the results of analyses from large amounts of data (Yamada *et al*., 2013; Delagenière *et al*., 2011; Fisher *et al*., 2015).

Recently, structure determination using multiple crystals has been facilitated by various developments that include the automation and acceleration of data collection. In structure determination using microcrystals, especially from lipid cubic phase (LCP) crystals of membrane proteins, structures are normally determined using the small-wedge synchrotron crystallography (SWSX) approach. In the SWSX approach, the total oscillation range is reduced, but the number of photons per oscillation width is increased. Since each wedge dataset covers only a part of the reciprocal space, tens to hundreds of datasets must be measured from many microcrystals that are mounted in different orientations (Cherezov *et al*., 2007; Rosenbaum *et al*., 2007). Finally, the acquired datasets are merged and used to determine the structure. In the case of serial femtosecond crystallography (SFX) (Barends *et al*., 2022) or serial synchrotron rotation crystallography (SSROX) (Gati *et al*., 2014; Hasegawa *et al*., 2017), a much larger number of images is required because every frame covers only a small portion of reciprocal space, posing challenges in data collection and analysis. Here, the automation of data collection and analysis is crucial and has provided opportunities for expanding the target and achieving structure determination for diverse protein samples (Healey *et al*., 2021).

During structural analysis using multiple crystals, it is crucial to select datasets that are sufficiently isomorphous. For example, if non-isomorphous datasets are merged in SAD phasing, it greatly complicates searching for the heavy atom positions (Giordano *et al*., 2012; Baba *et al*., 2021). Hierarchical clustering analysis (HCA) is an approach that has been successfully utilized for extracting highly isomorphous datasets from multiple crystals. This approach has been implemented in the program *BLEND* (Foadi *et al*., 2013) that conducts unit cell-based HCA, while HCA based on the correlation of diffraction intensities (also referred to as ‘intensity-based HCA’) has been realized in the program *ccCluster* (Santoni *et al*., 2017). The automatic data processing pipeline *KAMO* implements both types of HCA.

Even when a structure can be solved using a single crystal, there are benefits to using multiple crystals for analyzing the structure. One benefit is in terms of attainable resolution. The resolution of a given structure can be improved as the number of datasets for merging increases, because the signal-to-noise ratio of weak diffraction spots is improved. Another benefit is linked to the analysis of polymorphs. Physiologically meaningful structural polymorphs have been found by classifying diffraction data from many crystals. Most earlier studies used HCA to extract highly isomorphic data for single structure determination. However, some recent studies have demonstrated that HCA can be a powerful tool to classify structural polymorphs (Nguyen *et al*., 2022; Soares *et al*., 2022). In principle, HCA does not require any prior information on how many data clusters (*i.e.*, number of ‘polymorphs’ in the context of MX) are involved in the entire dataset. In practice, however, there are generally two ways to interpret the results from the HCA. One approach determines a threshold for the degree of ’isomorphism’ and clustered data below the threshold are considered to underly the same structure. The other approach decides the number of data clusters prior to analysis. Since it is impossible to know how many polymorphs are involved in the entire dataset, the former approach should be desirable. However, since there is no reference for deciding an appropriate threshold (‘isomorphic threshold’) at present, it is necessary to analyze the merged datasets at each cluster exhaustively, and the results should be interpreted case-by-case. Therefore, we investigated the feasibility of assigning an ‘isomorphic threshold’, which can be used to select the candidate data clusters where polymorphs may be identified.

Furthermore, we also investigated the usefulness of applying HCA to data collected by the helical scheme for capturing both inter and intra-crystal structural polymorphs. The helical scheme, strictly referred to as a ‘continuous helical scheme,’ was originally developed as a data collection scheme to avoid severe radiation damage (Flot *et al*., 2010). In contrast to the conventional single-point oscillation scan, crystals are translated during the data collection by the helical scheme. Therefore, the dose for each crystal volume can be considered constant. Since each frame is acquired from a different point in the crystal, splitting the full dataset into several partial datasets (chunks) could be useful to observe structural differences present in the crystal by processing each chunk individually and classifying them by HCA. Indeed, by using this approach, structure determination was possible even from heterogeneous crystals (Katoh *et al*., 2020).

In this paper, we sought to determine the ‘isomorphic threshold’ using *in silico* mixed datasets consisting of two different high-resolution datasets from standard test protein samples (trypsin). This threshold was then applied to two representative protein samples (nuclear transport receptor Transportin-1 and [NiFe]-hydrogenase maturation factor HypD) to evaluate whether the suggested threshold suitably classifies polymorphs.

## 2. Materials and Methods

### 2.1. Preparation of apo-and inhibitor-bound trypsin crystals

Bovine pancreas trypsin (Molecular weight: approximately 24 kDa, Fujifilm Wako Pure Chemicals) was dissolved in 25 m*M* HEPES pH 7.0 with 5 m*M* CaCl_2_ to a concentration of 30 mg ml^-1^. The precipitant solution was 30% (*w/v*) PEG 3350, 0.1 *M* Tris-HCl pH 8.5, 0.2 *M* Li_2_SO_4_. Crystallization was performed at 293 K by the sitting-drop vapor diffusion method using MRC-II plates (SWISSCI), and 200 μm crystals appeared within a few days. Several crystals were harvested before adding the compound to obtain data for apo-form. The crystals were soaked in cryoprotectant, which contained 10% (*v/v*) ethylene glycol mixed with a crystallization buffer, then cryocooled in liquid nitrogen (Yamane *et al*., 2011).

The inhibitor compound was directly added to the droplet on the crystallization plates using an Echo 650 acoustic liquid handler (Beckman Coulter). In this study, the following two inhibitors were used (Fig. S1): 4-Methoxybenzamidine and 5-Chlorotryptamine (hereinafter referred to as ‘benzamidine’ and ‘tryptamine’, respectively). Each inhibitor was added to the crystallization droplet at a final concentration of 10 m*M* containing 10% (*v/v*) dimethyl sulfoxide (DMSO). After addition of the inhibitor, the crystallization plate was placed at 293 K for an hour to allow sufficient inhibitor diffusion into the crystals. Inhibitor-bound trypsin crystals were fished from the crystallization plate, cryoprotected in a similar manner to apo-trypsin, and cryocooled in liquid nitrogen. DMSO concentration and the incubation time were determined based on the results of a preliminary study. In brief, several series of crystals with different DMSO concentrations and incubation times were prepared using apo-trypsin. Then, data were collected from these crystals to identify the conditions where the diffraction quality was not significantly degraded.

### 2.2. Diffraction data collection, data processing, and structure determination of trypsin

All diffraction data of trypsin crystals were collected at BL32XU, SPring-8, using an automated data collection system *ZOO* (Hirata *et al*., 2019). Data were obtained from four crystals for apo-form, benzamidine-bound, and tryptamine-bound trypsin crystals, respectively. All datasets were acquired using a continuous helical scheme for 360° oscillation with the following experimental parameters; Oscillation width: 0.1°, Exposure time 0.02 s., Beam size: 10 μm [H] × 15 μm [V], Wavelength: 1 Å, Average dose/crystal volume: 10 MGy, Detector: EIGER X 9M (DECTRIS Co. Ltd.). *KUMA* module in *ZOO* automatically estimated the attenuation factor from measured crystal size and designated dose value (Hirata et al., 2019). The rotation axis of the goniometer is horizontal to the ground. Basically, the irradiation vector was set in the apparent long-axis direction of the crystal, which roughly coincided with the rotation axis, in the ZOO helical scheme.

The obtained data were automatically processed by *XDS* (Kabsch, 2010) exploited in *KAMO* (Yamashita *et al*., 2018). Subsequently, automated structural analysis was performed using *NABE* (Matsuura *et al*., unpublished), an automated structural analysis pipeline currently under development at SPring-8 MX-BLs. *NABE* provides an interface to manage all the merged data in clustering results generated by *KAMO* by summarizing the data statistics and the electron density map. *NABE* runs *DIMPLE* (Wojdyr *et al*., 2013) on the data processed by *KAMO*, using the given template model for molecular replacement. If the amino acid residues and atom names are specified for a site of interest, *NABE* automatically generates pictures of the protein model and resultant electron density maps (2*F*_o_-*F*_c_ and *F*_o_-*F*_c_ maps) around the selected site by using *Coot* (Emsley & Cowtan, 2004), *Raster3D* (Merritt & Bacon, 1997), and *ImageMagick* (The ImageMagick Development Team, 2021; Available at https://imagemagick.org). The pictures are not a static snapshot but a GIF image that rotates, making it easier to see the surrounding environment, including the depth direction, which is difficult to perceive from a static image. After the above processes are completed, *NABE* returns a HTML report that tabulates the resolution of the obtained data, *R*_Free_, *B*-factor, etc., from the results of *KAMO* and *DIMPLE*, along with the GIF image for each dataset. When multiple datasets are used, for example during SWSX measurements, *KAMO* classifies the diffraction data by HCA and then merges them in multiple clusters. Subsequently, *NABE* automatically performs the instantaneous data analysis for all these merged datasets.

Here, using the report from *NABE*, we listed the electron density map around the inhibitor binding site of trypsin and confirmed that the *F*_o_-*F*_c_ density for inhibitors was not observed in the apo-trypsin. In contrast, *F*_o_-*F*_c_ density for each inhibitor was observed for benzamidine-bound and tryptamine-bound trypsin datasets (Fig. S1).

### 2.3. Clustering analysis using data from apo-and inhibitor-bound trypsin

In HCA, an ‘isomorphism’ between the datasets is represented as a vertical ‘distance’ in the dendrogram. Isomorphous datasets are linked within smaller distances, while other, more distantly related datasets are linked with longer distances. Supposing there are several structural polymorphs in the multiple datasets, each polymorph will make a cluster consisting of datasets within a certain ‘threshold’ (referred to as ‘isomorphic threshold’) in the dendrogram. In actual data, the number of polymorphs involved in the obtained dataset is unpredictable. Therefore, two apparently different datasets were used to investigate an ‘isomorphic threshold’ for identifying polymorphs. For this purpose, we used high-resolution datasets from apo-and two different inhibitor-bound trypsin.

Two different parameter-based HCAs are implemented in *KAMO*: one is unit cell-based, and the other is intensity-based HCA. *KAMO* performs unit cell-based HCA using *BLEND*. Intensity-based HCA is performed based on the CC calculation by the *cctbx* (Grosse-Kunstleve *et al*., 2002) method miller_array.correlation.coefficient and grouping by the *scipy* (Virtanen *et al*., 2020) method cluster.hierarchy.dendrogram. The structural changes of protein molecules in the crystal possibly affect the unit cell constants or diffraction intensities. These two different parameter-based HCA are somewhat different in terms of isomorphism. In the unit cell-based HCA, the classification is based on the isomorphism of the cell parameters, which reflects a more macroscopic aspect rather than the protein structures. In contrast, intensity-based HCA can detect much smaller structural changes.

Here, we examined how the two different datasets are classified by two different parameter-based HCAs: unit cell-based and intensity-based. To evaluate whether splitting the data collected by the helical scheme enables polymorph analysis within the same crystals, collected data were divided into 30° chunks and individually processed using *KAMO* with the split_data_by_deg=30.0 option. The test dataset for clustering analysis was prepared by mixing the same number of diffraction datasets from two different structures. In this study, 360° data collected by the helical scheme were used with division into 30° chunks. Here we used forty-eight 30° chunks of each structure, corresponding to datasets from four crystals. Two parameter-based HCAs were applied to the *in silico* mixed datasets consisting of 96 chunks from the following combinations of two datasets (48 + 48 chunks): (1) apo-and benzamidine-bound trypsin, and (2) benzamidine-bound and tryptamine-bound trypsin. HCA and data merging were carried out using *KAMO* (*kamo.auto_multi_merge*) in the following scheme. Prior to HCA, *KAMO* selects the datasets using for the merging step (referred to as ‘pre-processing’). Firstly, the equivalent data group is selected based on P1 symmetry. Then, the selected datasets are filtered based on the unit cell constant using Tukey’s criterion. HCA is performed for the filtered data list, and merged data will be generated at each cluster. KAMO rejects data on a frame-by-frame and dataset-by-dataset basis based on crystallographic statistics in the three cycles of data merging. The electron density map for the merged data at each cluster was depicted to evaluate the effect of data contamination. Molecular replacement was performed with the template model (PDB ID: 3RXA). Evaluation of the electron density map was performed using *NABE*.

In general, the definition of ‘isomorphism’ between datasets and the linkage method to calculate the distance between clusters contribute significantly to the resultant dendrogram from HCA. For intensity-based HCA, correlation coefficient (CC) of intensity is used as an indicator of isomorphism. A different definition of distance is used in *KAMO* and *ccCluster* as a default setting; [1-CC]^1/2^ is used in *KAMO* (Yamashita *et al*., 2018), while [1-CC^2^]^1/2^ is used in *ccCluster* (Santoni *et al*., 2017). In this study, we used [1-CC^2^]^1/2^ (hereafter referred to as *d*_CC_) based on the preliminary investigations on the following distance definitions available in *KAMO*: 1-CC, [1-CC]^1/2^, and [1-CC^2^]^1/2^. For the linkage method, the ‘Ward’ linkage is often used, as it is empirically less prone to causing the ‘chain effects’ and ‘inversions’ of the dendrogram (Murtagh & Legendre, 2014). The program *BLEND* also adopts the Ward method for the same reason (Foadi *et al*., 2013), while the program *ccCluster* uses the ‘average’ linkage method instead. In intensity-based HCA implemented in *KAMO,* the Ward linkage is utilized as the default setting. Based on the results with seven linkage methods available in *scipy* module (Section S1 and Fig. S2), we used the Ward linkage in this study. In the Ward linkage, the closest cluster is selected by minimizing the increase of the variance when joining the clusters. When merging a pair of single datasets, the Ward distance can be considered as synonymous with the *d*_CC_, so that the CC value can be calculated directly from the value of Ward distance. Whereas, when merging clusters containing multiple datasets, the distance between these clusters is determined based on the data contained in the cluster. Accordingly, the Ward distance does not directly give the CC value for each dataset inside the clusters. For example, Ward distance = 0.6 does not mean that CC value for each pair of content dataset equals to 0.8.

### 2.4. Determination of isomorphic threshold from observed/simulated data

Based on the results of HCA of trypsin *in silico* mixes, we hypothesized that the presence of polymorphic structures could be detected by the Ward distance threshold, on the vertical axis of the dendrogram. If there are multiple clusters below this "isomorphic threshold," each cluster is considered as different structures. However, the absolute values of Ward distance generally increase with the number of datasets, and they also increase when the CC distributions among different structures are diverse. Therefore, it would not be useful to derive the absolute Ward distance threshold directly from the dendrogram of observed trypsin data as a general-purpose index. Therefore, we simply define the isomorphic threshold using the maximum Ward distance (W_0_ in Fig. S3) of the entire system as follows.

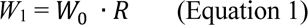

(W_0_: maximum Ward distance, W_1_: isomorphic threshold, R: a constant to be determined here, a ratio with magnitude between 0-1)

The "isomorphic threshold" is defined as the larger Ward distance between two branches when two structures can be classified (W_1_ in Fig. S3). To determine R in Equation 1, the simulation was performed with the following steps. Based on the apo-and benzamidine-bound trypsin datasets, we firstly modeled the intensity CC distribution for all of three combinations. Assuming that there are multiple datasets and two structures are included, we created the CC matrix for HCA using CCs those follow the probability density of our model (Fig. S3). We performed HCA from the CC matrix and evaluated whether the two structures could be classified as their original labels on the dendrogram.

Firstly, we checked whether the simulation successfully reproduced the dendrograms obtained experimentally. Since W_0_ and W_1_ can be obtained from the dendrogram in the HCA simulation, they were simply compared to the observed values. Next, the CC model parameters were changed from the initial model to make classification increasingly difficult, and calculations were performed stepwise until the final condition where classification was no longer possible. The number of datasets was varied from 100 to 1000, each with half different two structural labels. The R calculated from W_1_/W_0_ and the score in each HCA were plotted for each parameter to determine suitable R to calculate the isomorphic threshold in the intensity-based HCA. The details of the simulation are described in section S2.

### 2.5. Application to representative sample 1: Polymorph analysis of nuclear transport receptor Transportin-1 in complex with a nuclear localization signal peptide

To evaluate whether the ‘isomorphic threshold’ suggested from the investigations on trypsin datasets is effective, HCA was carried out in representative sample datasets with the proposed ‘isomorphic threshold’.

Transportin-1 (Trn1) is one of the nuclear transport receptors which recognizes a nuclear localization signal (NLS) sequence harbored in cargo proteins and brings them into the nucleus. Detailed information about sample preparation of Trn1-NLS peptide complex (molecular weight is approximately 98 kDa and 2.5 kDa, respectively) is described in section S3. Here we describe it briefly. The Trn-1 Δloop mutant was produced using an *Escherichia coli* expression system. Purification of Trn1 was performed using Glutathione Sepharose and anion exchange columns followed by size exclusion chromatography. NLS peptide of Trn1 (Eurofins) was dissolved in the purification buffer (110 m*M* potassium acetate, 200 m*M* HEPES-KOH, 10 m*M* DTT). Trn1-NLS peptide complex (hereinafter referred to as ‘Trn1-peptide complex’) was prepared by mixing 5 mg ml^-1^ Trn1Δloop and 5 m*M* NLS peptide. Trn1-peptide complex crystals were obtained under the crystallization condition containing 0.5 *M* NaK phosphate pH 5.0. The obtained crystals were cryo-cooled in liquid nitrogen after cryoprotection with a 30% (*w/v*) glycerol-containing reservoir solution. Diffraction data were automatically collected at BL32XU, SPring-8 using *ZOO*. From each of the four crystals, 720° rotation data were acquired by the continuous helical scheme. Other experimental parameters were identical as performed for the trypsin crystals: Oscillation width: 0.1°, Exposure time 0.02 s., Beam size: 10 μm [H] × 15 μm [V], Wavelength: 1 Å, Average dose/crystal volume: 10 MGy, Detector: EIGER X 9M (DECTRIS Co. Ltd.). *KUMA* module in *ZOO* automatically estimated the attenuation factor from measured crystal size and designated dose value. The obtained 720° data was split into 30° chunks, and hierarchical clustering was applied. The electron density map was compared using *NABE* with the peptide-free structure model as a template model for MR. The peptide-free model was prepared prior to the clustering analysis by the following procedures: phasing by molecular replacement using the program *PHASER* (McCoy *et al*., 2007) with a template model (PDB ID: 5YVI), followed by an iteration of refinement using *phenix.refine* (Liebschner *et al*., 2019) and manual model building using *Coot*.

### 2.6. Application to representative sample 2: Polymorph analysis of [NiFe]-hydrogenase maturation factor HypD

HypD is one of the maturation factors of [NiFe]-hydrogenase and can form a complex with other maturation factors (Muraki *et al*., 2019). The C360S variant of HypD from *Aquifex aeolicus* (hereinafter referred to as *Aa*HypD-C360S, molecular weight: approximately 42 kDa) was produced using an *Escherichia coli* expression system and purified using a cation exchange column with a sodium chloride gradient and size exclusion chromatography. *Aa*HypD-C360S crystals were obtained 16% (*w/v*) polyethylene glycol 3350, 0.1 *M* citrate buffer pH 5.6, and 1 m*M* dithiothreitol as a reservoir solution. The obtained crystals were cryo-cooled in liquid nitrogen after cryoprotection. Diffraction data were automatically collected at BL45XU, SPring-8 using *ZOO*. From each of the six crystals, 360° rotation data were acquired by a continuous helical scheme. Other experimental parameters are listed as follows; Oscillation width: 0.1°, Exposure time 0.02 s., Detector: PILATUS3 6M (DECTRIS Co. Ltd.). *KUMA* module in *ZOO* automatically estimated the attenuation factor from measured crystal size and designated dose value. The obtained 360° data was split into 30° chunks, and hierarchical clustering was applied. Data were merged at largely separated data clusters that should exhibit structural polymorphs. Evaluations of the electron density map were carried out by *NABE*. Before the analysis by *NABE*, a template model was prepared using *PHASER* (PDB ID for the initial model: 2Z1D), *REFMAC5* (Murshudov *et al*., 2011), and *Coot*.

## 3. Results and Discussions

### 3.1. Test study: Classification of different trypsin datasets

To investigate whether HCA can classify polymorphic datasets, *in silico* mixed datasets were prepared, which consist of 48 chunks each from two different (apo-and benzamidine-bound) trypsin datasets resulting in a total of 96 chunks submitted to *KAMO*. The resolution of each chunk was approximately 1.2 Å. Each chunk contained about 50,000 reflections, and the overall completeness was about 40% and the space group was *P*2_1_2_1_2_1_. *KAMO* rejected 13 chunks during pre-processing as described in the method section leaving 83 chunks for HCA (Table S1). Reflections up to 1.50 Å were used to calculate the correlation between the datasets in the intensity-based HCA.

#### 3.1.1. Classification of diffraction data for apo-trypsin and benzamidine-bound trypsin

Unit cell-based and intensity-based HCAs were applied to the *in silico* mixed datasets, including apo-and inhibitor-bound trypsin (Figs. 1 and 2). In both HCA results, merged data for the top cluster (Cluster 82) exhibited no significant *F*_o_-*F*_c_ density corresponding to benzamidine at 3.0σ, even though chunks of benzamidine-bound trypsin datasets were included in the mixed dataset (Table S2). This may be explained by the fact that the number of apo-trypsin datasets in the merged data was larger than that of benzamidine-bound trypsin as a consequence of pre-processing by *KAMO* before HCA.

**Figure 1.**
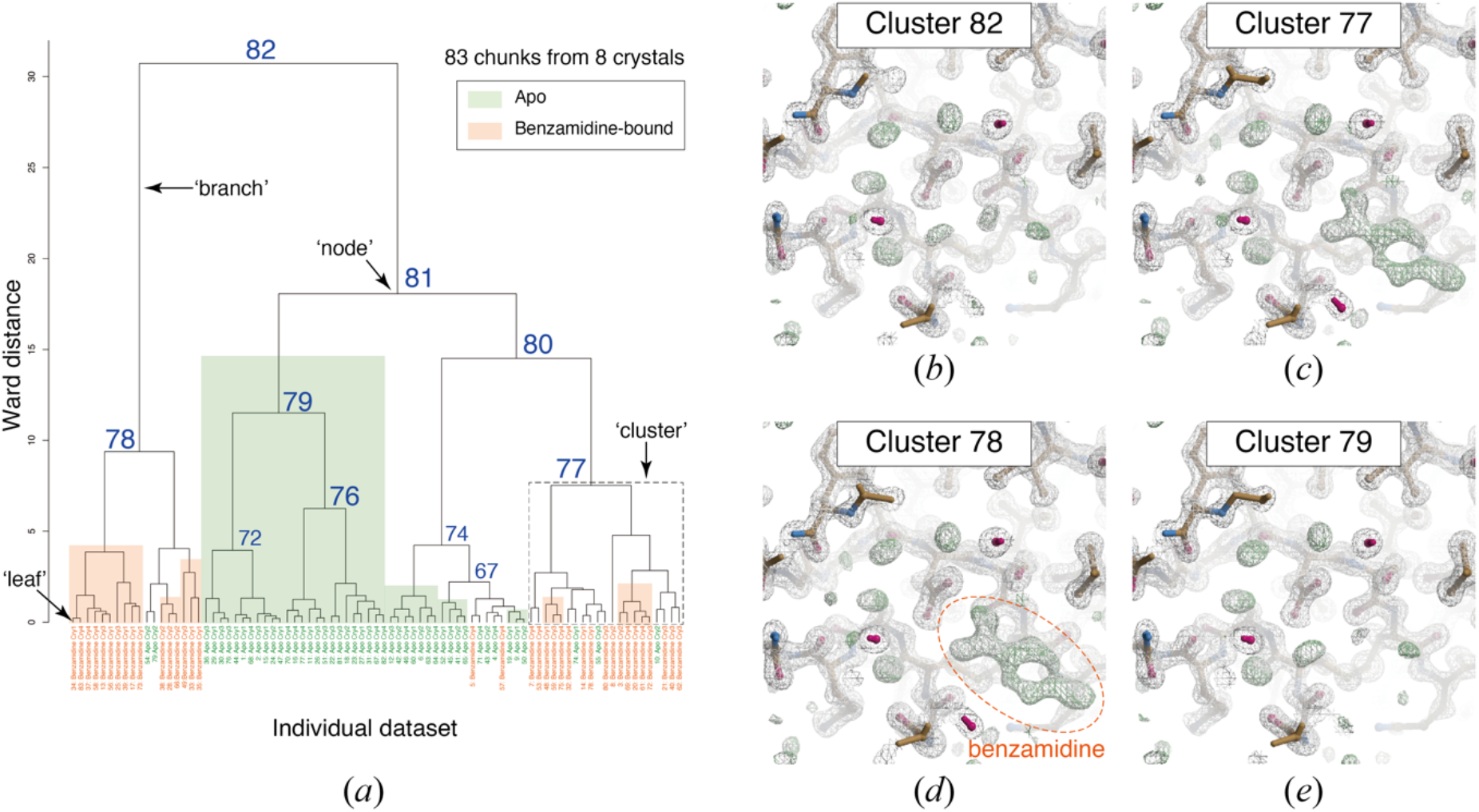
Result of unit cell-based HCA of *in silico* mixed datasets containing apo-trypsin and benzamidine-bound trypsin. (a) Resultant dendrogram. The definition of ‘node’, ‘branch’, and ‘cluster’ used in this study is illustrated in the dendrogram. Cluster numbers and data labels are depicted on the resultant dendrogram of unit cell-based HCA. The blue numbers indicate cluster numbers at each node. Chunks for apo-trypsin are shaded in green, and those for benzamidine-bound trypsin are colored in orange. (b)-(e) Electron density map around the inhibitor binding site of trypsin in different clusters. Contour level of 2*F*_o_-*F*_c_ map (gray mesh) is 1.0σ, and that of *F*_o_-*F*_c_ map (green mesh) is 3.0σ. Maps were generated by *Coot* exploited in the *NABE* system. See also Fig. S2.

**Figure 2.**
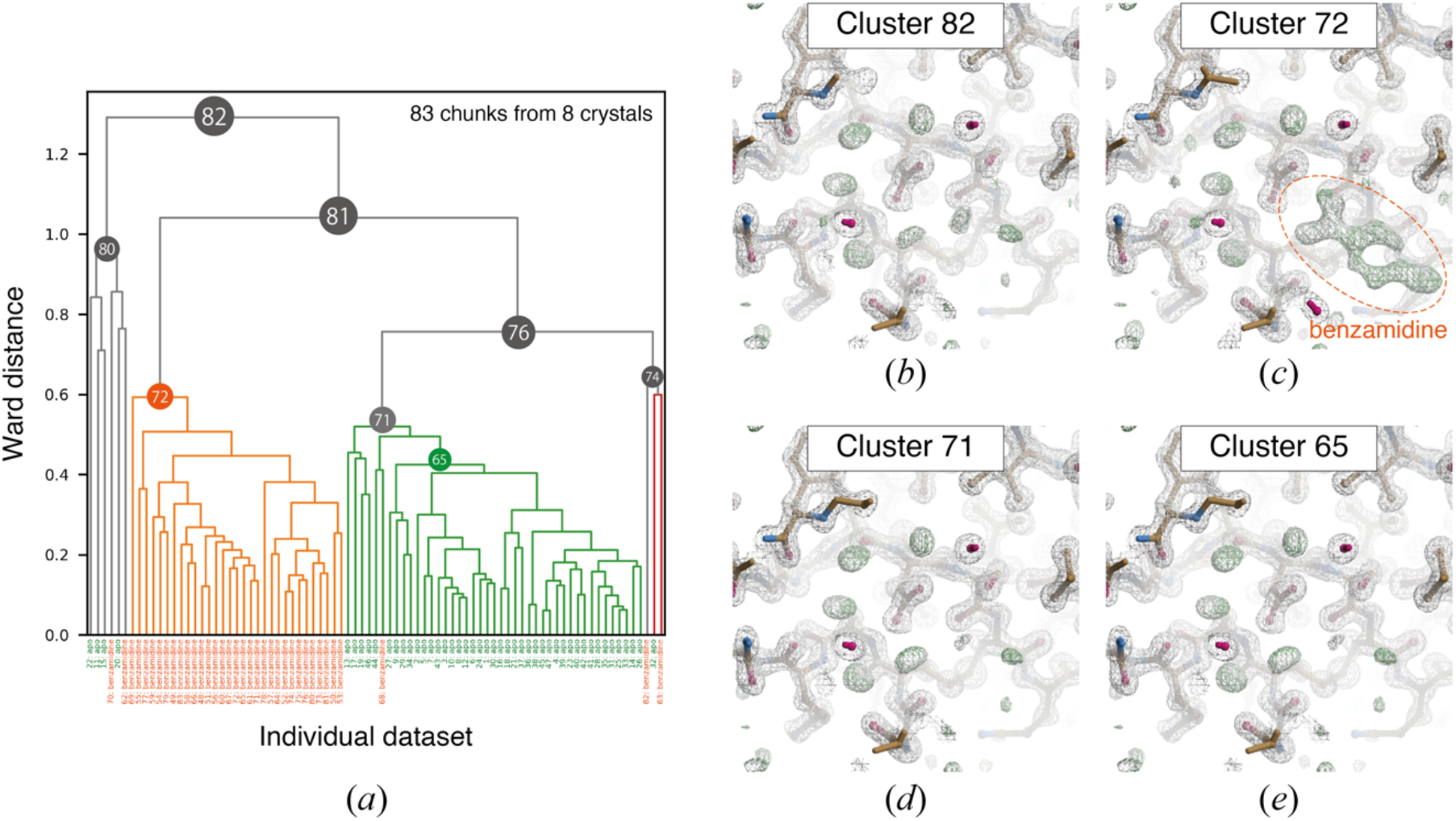
Result of intensity-based HCA on *in silico* mixed dataset containing apo-trypsin and benzamidine-bound trypsin. (a) Resultant dendrogram. Data labels are shown at the bottom: (green) apo-trypsin and (orange) benzamidine-bound trypsin. The color threshold for the dendrogram is set to 0.6. (b)-(e) Electron density map around the inhibitor-binding site obtained from the merged data in different clusters. Contour levels of 2*F*_o_-*F*_c_ map (gray mesh) and *F*_o_-*F*_c_ map (green mesh) are 1.0α and 3.0α, respectively. Maps were generated by *Coot* exploited in the *NABE* system.

The largest linear cell variation (LCV) value (Foadi *et al*., 2013), a characteristic quantity for unit cell variation, was 0.59%. Due to the slight differences in the unit cell constant between apo-trypsin and benzamidine-bound trypsin (Fig. S4), unit cell-based HCA did not succeed in fully separating apo-trypsin and benzamidine-bound trypsin datasets (Fig. 1). This implies that the unit cell-based HCA is less sensitive to a small lattice change like the one found between apo-trypsin and benzamidine-bound trypsin crystal forms. In practical cases, well-classified clusters should be inferred from the electron density map since there is no label for polymorphs. Accordingly, we investigated the electron density map obtained from merged data at each cluster. The two different datasets appeared to be separated at the first branch of the dendrogram (Clusters 78 and 81) based on the electron density map. Although both the apo-and benzamidine-bound chunks are involved in these clusters, data contamination cannot be distinguished from the electron density map. Due to the rejection of outlier datasets during the merging process by *KAMO*, some of the contamination data chunks were eliminated. For example, although Cluster 78 has 2 apo-trypsin and 16 benzamidine-bound trypsin chunks immediately after the clustering, 10 chunks containing only benzamidine-bound trypsin datasets were merged as the final dataset resulting in a clear benzamidine electron density to be present (Table S2 and Fig. 1). In most of the other cases, the final merged data still contained some contaminations even after the outlier rejection (Table S2). For instance, the merged data at Cluster 77 consisted of 2 apo-trypsin and 11 benzamidine-bound trypsin chunks. However, a small number of contaminations did not affect the resulting electron density map (Fig. 1). Approximately 15% of the contamination did not affect the electron density map resulting from the dominant dataset in this test case.

In contrast to the above-described results, the intensity-based HCA (Fig. 2) succeeded to classify the two mixed datasets completely (Clusters 71 and 72). Clear *F*_o_-*F*_c_ density for benzamidine was observed at Cluster 72, while no significant *F*_o_-*F*_c_ density of benzamidine was observed in other clusters at 3.0σ. Cluster 71 had only one benzamidine-bound chunk, and there may be two possibilities why this chunk was involved in the apo-trypsin cluster: either the occupancy of benzamidine was low or the data quality was poor. In consideration of the former possibility, the data obtained from a selected benzamidine-bound crystal was divided into four 90° chunks and performed occupancy refinement of benzamidine with *REFMAC5*. As a result, occupancies of 97%, 93%, 97% and 92% were not significantly different, suggesting that the ligand occupancy was likely to be constant in the whole crystal volume. With regard to the latter possibility, automated helical data collection on large crystals often results in significantly low diffraction power at both ends of the crystal. This point is discussed again in section 3.1.4.

In the intensity-based HCA, CC is the key information that has an influence on the results of classification. In CC calculation, common reflections up to the specified resolution are used. To investigate the resolution dependency on CC distance (*d*_CC_) value, intensity-based HCA was performed at several resolution cut-off (Fig. S5). Despite lowering the cut-off resolution to 3.5 Å for CC calculation, intensity-based HCA successfully classified apo-and benzamidine-bound trypsin data chunks. Accordingly, resolution dependency on the CC calculation appeared insignificant as far as we investigated. The successful sorting of apo-and benzamidine-bound data chunks demonstrates that intensity-based HCA can be effective to disentangle heterogeneous data chunks, even if differences in cell dimensions are too minute to be classified by unit cell-based HCA.

#### 3.1.2. Classification of two different inhibitor-bound trypsin datasets

Next, we tested HCA-based classification on datasets obtained from crystals of trypsin with two different inhibitors that have different skeletal formulas (Fig. S1) but share the same binding site; benzamidine and tryptamine.

Although clusters of homogenous datasets can be found for some clusters (*e.g.,* Clusters 74 and 76), the two different inhibitor-bound data could not be successfully sorted by unit cell-based HCA (Fig. 3). The largest LCV value was 0.75%, slightly larger than the case for apo-and benzamidine-bound trypsin. This implies that a larger LCV value of 0.75% is still insufficient to classify the different trypsin datasets using the unit cell-based HCA. Based on the electron density map (Fig. S6), the two different datasets appeared to be roughly separated at the first branch (Clusters 81 and 82). However, the electron density map at Cluster 80 clearly exhibited a mixture of both types of inhibitor-bound data. It appears to be difficult to distinguish the base skeletal structure from the electron density map. The merged data consisted of 6 benzamidine-bound chunks and 14 tryptamine-bound chunks, resulting in approximately 30% contamination against the dominant datasets (Table S3). At Cluster 77, the merged data consisted of 9 benzamidine-bound chunks and 4 tryptamine-bound chunks, where about 31% of contamination was against the prevailing dataset. Based on these results, approximately 30% contamination by a minor dataset might be a tolerable upper limit for obtaining an electron density map that allows initial model building.

**Figure 3.**
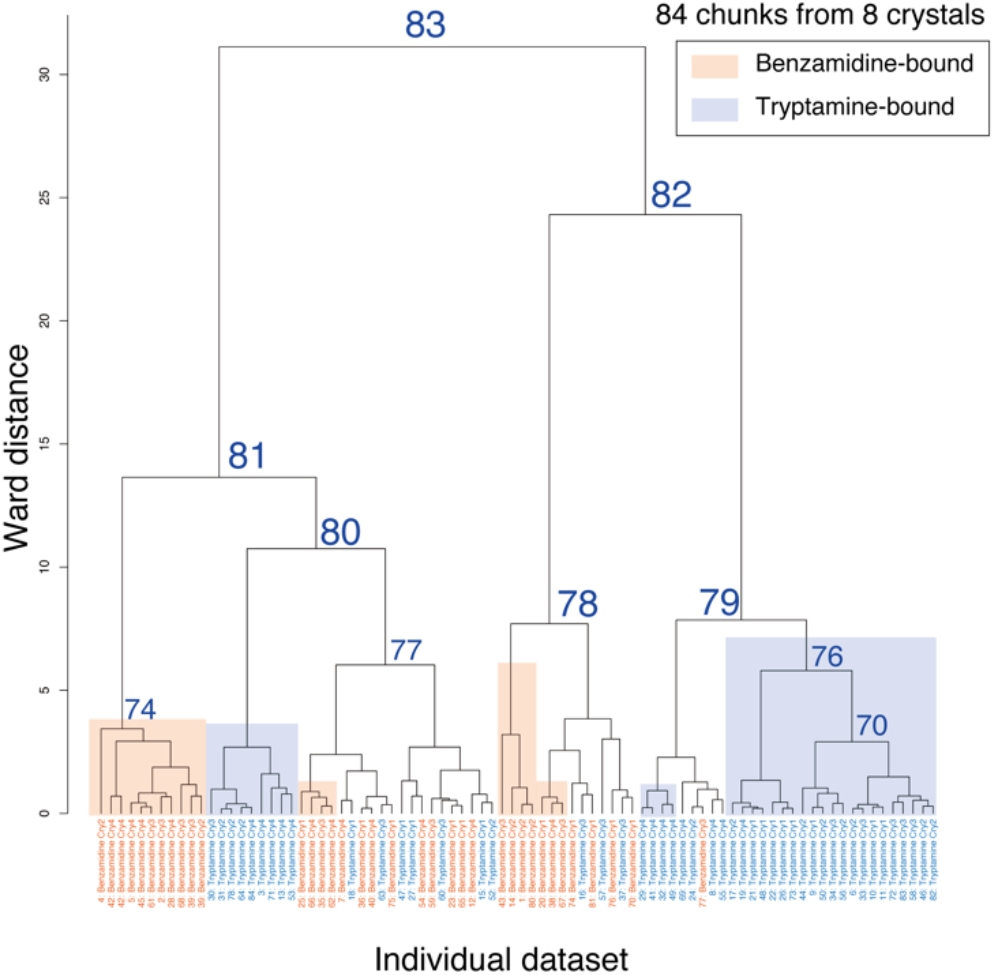
Dendrogram from unit cell-based clustering of mixed datasets including benzamidine-bound and tryptamine-bound trypsin. Cluster numbers and data labels are depicted on the resultant dendrogram of unit cell-based HCA. To evaluate the clustering effect, each cluster leaf is shaded according to the original dataset: (orange) benzamidine-bound and (blue) tryptamine-bound trypsin. The number labeled at each cluster node indicates the cluster number output from BLEND.

Intensity-based HCA (Fig. 4) successfully sorted the data chunks into two homogenous datasets (Clusters 72 and 74), although two inhibitor-bound datasets were mixed at some small clusters in the right branch (Clusters 73, 78, and 81). The electron density maps obtained at Clusters 74 and 72 exhibited the clear *F*_o_-*F*_c_ densities derived from benzamidine and tryptamine at 3.0σ, respectively (Fig. S7).

**Figure 4.**
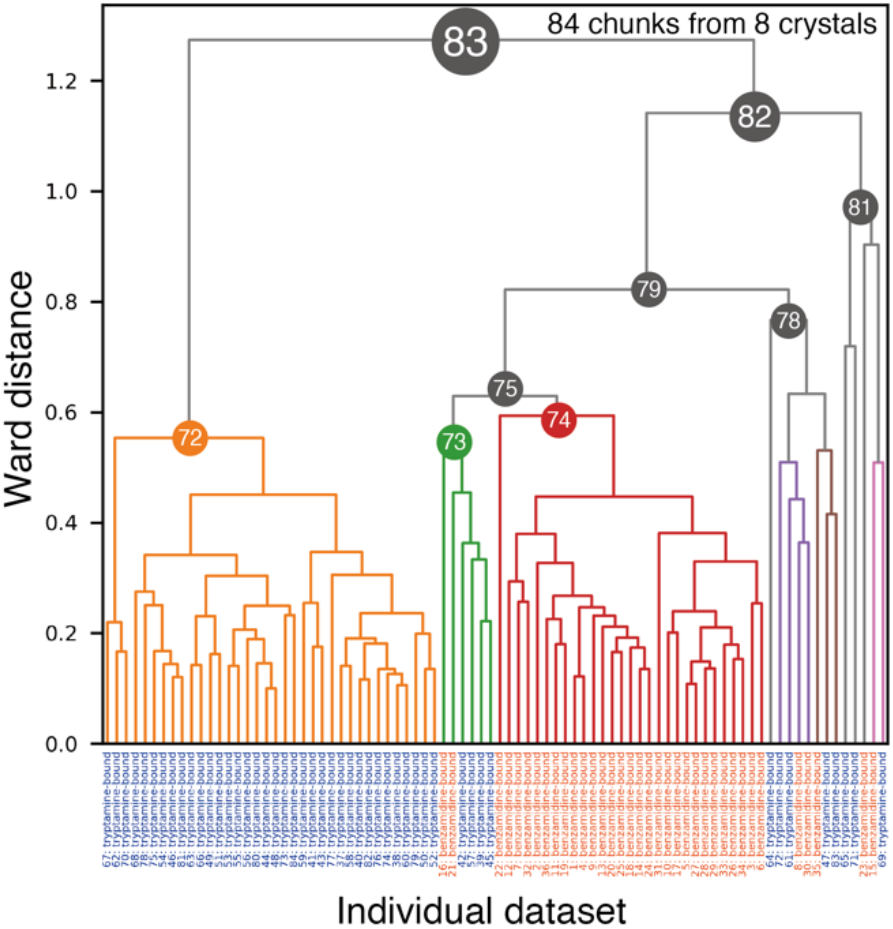
Dendrogram from intensity-based HCA of mixed datasets including benzamidine-bound and tryptamine-bound trypsin. Data labels at the bottom are (orange) benzamidine-bound and (blue) tryptamine-bound trypsin.

#### 3.1.3. Classification of apo-trypsin and two inhibitor-bound trypsin datasets

As shown in the earlier sections, two different datasets were classified using a single-step intensity-based HCA. However, more than three structural polymorphs may be found in practical cases. For example, when clustering is applied to data obtained in a time-resolved experiment, there may be more than three intermediates before and after the intermediate of interest. To test such a case, intensity-based HCA was applied to an *in silico* mixed datasets containing all three different trypsin datasets used in this study, namely apo-trypsin, benzamidine-bound, and tryptamine-bound trypsin datasets.

The results showed that intensity-based HCA classified three different datasets almost perfectly (Fig. S8), while the unit cell-based HCA did not classify these datasets well (Fig. S9). At the first branch in Figure S8, one cluster contains only tryptamine-bound chunks (Cluster 115) and the other mainly consists of apo-and benzamidine-bound chunks (Cluster 130). From the further clustering of the latter cluster, the homogeneous data cluster for apo-trypsin appeared at Cluster 114 and that for benzamidine-bound trypsin appeared at Cluster 116. During the separation, some small clusters (Clusters 119, 120, and 128) appeared. These clusters are considered outliers because the number of chunks involved is less, and the distances between the chunks are relatively large compared to the other clusters. Although some inhibitor-bound chunks were mixed at the apo-trypsin dominant cluster (Cluster 114), it is possibly due to lower inhibitor occupancies or relatively low-quality data.

#### 3.1.4. ‘Isomorphic threshold’ suggested from the investigations on trypsin observed datasets

CC values for the datasets in the clusters for the in silico mixed trypsin (apo-and benzamidine-bound trypsin; section 3.1.1) were 0.93 and 0.94 for the clusters of apo-trypsin (Cluster 71) and benzamidine-bound trypsin (Cluster 72) datasets, respectively. The result suggests that HCA is effective in the classification of such small structural changes. Therefore, we investigated the change of CC by a structural change of a certain volume of protein moiety. The change of CC value was examined for rotation of the partial or whole trypsin molecule (223 amino acids (AA) comprises the full length) without any changes in the unit cell constants (Fig. S10). It was found that even a relatively significant conformational change of 5° rotation of the terminal 10 AA-long helix resulted in a CC decrease of 0.015 compared to the original structure. When a quarter (57 AA) of trypsin residues were rotated, a 5° rotation resulted in a CC change of approximately 0.030. The simulation also proved that larger CC changes occur if the whole molecule was rotated.

For the 30° chunks from apo-(referred to as ‘apo’) and benzamidine-bound (referred to as ‘benz’) trypsin used in the present study, a histogram of the *d_CC_* values obtained for each combination of chunks is illustrated in Figure S11. The distribution of *d_CC_* for homogeneous pairs of apo chunks was centered around 0.2, whilst for the *d_CC_* obtained between heterogeneous data of apo-and benz-trypsin had a distribution shifted slightly to the right, around 0.25 (CC ∼ 0.97).

The histograms showed that even among homogeneous data, some portions of the combination exhibited a high *d_CC_* value greater than 0.6 (Fig. S11). The heat map of *d_CC_* values between all pairs of datasets revealed that there are some datasets that do not have any correlation against almost all the data (Fig. S12). These data were mostly chunks emanating from the tip of the crystals. Since the selection of equivalent datasets and unit cell-based filtering are performed prior to clustering, these chunks are similar to the other chunks at least with respect to the cell parameters. The chunks at the tip of crystals did not significantly affect the structural analysis as discussed in the previous section, therefore, the calculation of CC may be not reliable (e.g., <*I*/α*I*> or the resolution limit for used intensities should be carefully considered during the CC calculation). However, all the data classified by HCA were included in the main distribution with a *d_CC_* of around 0.2, indicating that the main *d_CC_* distribution is important for accurate classification by HCA.

As described in section 2.4 and section S2, numerical simulations were performed to determine an ’isomorphic threshold’ from apo-and benz-trypsin datasets. The observed median values for CC_apo-apo_, CC_benz-benz_, and CC_apo-benz_ distributions were 0.978, 0.970, and 0.962, respectively, with corresponding standard deviations of 0.020, 0.019, and 0.017. Only data satisfying d_CC_ < 0.4 were used to characterize the prominent peak of the CC distribution for these statistics. Distributions of CCs and fitted curves are shown in Fig. S13(a).

The HCA simulations for apo-and benz-trypsin showed perfect classification without contamination on the dendrogram (Fig. S13(b)). After repeating 100 calculations, the mean and standard deviation of W_1_ for classifying the two different structures were 0.61 and 0.03, respectively. The result demonstrates that our simulation roughly reproduces the observed data.

In the next step, the validated model was used to perform HCA simulations by modifying its parameters to make it difficult to classify two structures. Figure 5 displays the relationship between the ratio R and HCA simulation scores. The lowest R in each plot, shown as a ’cross mark,’ is the result of HCA using the original model. From there, as the classification gets difficult, R increases, and the score worsens. The R greater than 0.7 worsens the classification score for all plots except the 1000 dataset. We regard the score threshold for success in classification as 0.9 (dotted line in Figure 10), then 0.6 to 0.7 can be a reasonable R for several hundred datasets. Based on these results, we assumed that the "isomorphic threshold" for polymorphism detection in intensity-based HCA can be calculated by multiplying W_0_ by 0.6-0.7.

**Figure 5.**
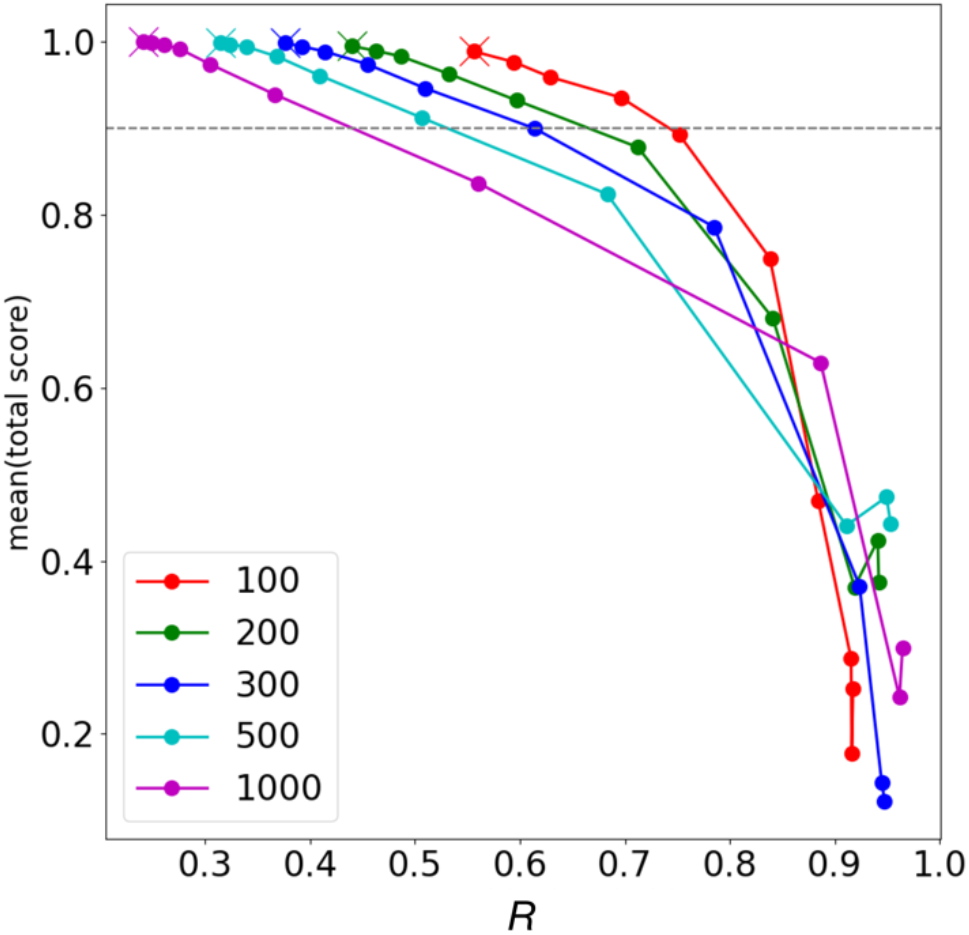
*R* in equation 1 and classification score in our HCA simulations (details are described in sections 2.4 and S2). Each line plot illustrates the scores assuming 100, 200, 300, 500, and 1000 datasets. The lowest *R* in each plot is shown as cross point and is the result of HCA using the original CC model (Fig S13(a)). HCA simulations were performed with the CC_apo-benz_ and CC_benz-benz_ positions gradually approaching each other in 10 steps, making classification more difficult. The rightmost plot shows the same *loc*s in CC_apo-benz_ and CC_benz-benz_.

By using the assumed ratio, classifications in the observed intensity-based HCA were examined. For the apo-/benz-trypsin case, 0.77∼0.90, the highest value of the dendrogram in Figure 2 (1.29) multiplied by 0.6∼0.7, was used as the isomorphic threshold. As a result, clusters 71, 72, and 74 were candidates for structural polymorphs, but 74 was not a complete dataset and could not be used for structural analysis. For two ligand-binding trypsins (Figure 4), the isomorphic threshold is 0.76∼0.89, which classifies clusters 72, 75, and 78 as candidates for structural polymorphs. Although cluster 73 is contaminated, cluster 75 shows a benzamidine-bound structure on the electron density map in Figure S7. A similar result is found in the intensity-based HCA on a mixed dataset, including apo-and benzamidine-bound trypsin. In Figure S8, the isomorphic threshold is about 0.96∼1.1, and clusters 115, 123, and 125 are candidates of polymorphs. Of these, clusters 115, 114, and 116 remain after excluding the obvious outlier clusters. Our isomorphic threshold proved to be a good indicator for classifying polymorphism in all trypsin cases.

The investigations on trypsin datasets indicated that intensity-based HCA can successfully classify structural polymorphs. The results also suggested that a single-step intensity-based HCA could be sufficient when the variation in unit cell constant is small, for example, when the largest LCV value is less than 1%. In the dendrogram from intensity-based HCA on trypsin test cases (Figs. 2, 4, and S8), the same datasets appeared to be clustered within our isomorphic threshold. Based on the results, polymorphs could be identified by grouping the datasets using a single-step of intensity-based HCA with the suggested ‘isomorphic threshold’. Since multiple steps of clustering (Nguyen *et al*., 2022) reduce the number of datasets in the final step by filtering the data during the clustering, fewer steps of clustering may be helpful when the total number of data is limited.

### 3.2. Applications to the representative samples

To evaluate whether the single-step of intensity-based HCA with our suggested ‘isomorphic threshold’ is useful for the detection of polymorphs in practical samples, we performed the intensity-based HCA and structure analysis on two representative examples: Trn1-peptide complex and *Aa*HypD-C360S. Here we selected the clusters that satisfy our threshold, alongside the completeness of the merged data being high enough to allow further structural analysis.

#### 3.2.1. Diffraction data classification and polymorph analysis of the Trn1-peptide complex

Diffraction data of the Trn1-peptide complex were obtained using a helical scheme for 720° rotation. Each dataset was divided into 30° chunks, yielding 96 chunks from 4 crystals. The resolution for each chunk was in the range of 3 to 4 Å. The number of reflections in each 30° chunk was approximately 60,000, the overall completeness was about 25% and the space group was *C*2. Reflections up to 3.67 Å were used to calculate the correlation between the datasets in the intensity-based HCA.

The obtained chunks were subjected to intensity-based HCA using *KAMO* (Fig. 6). Although some outliers were observed, two major clusters (Clusters 72 and 76) were selected for further structural analysis.

**Figure 6.**
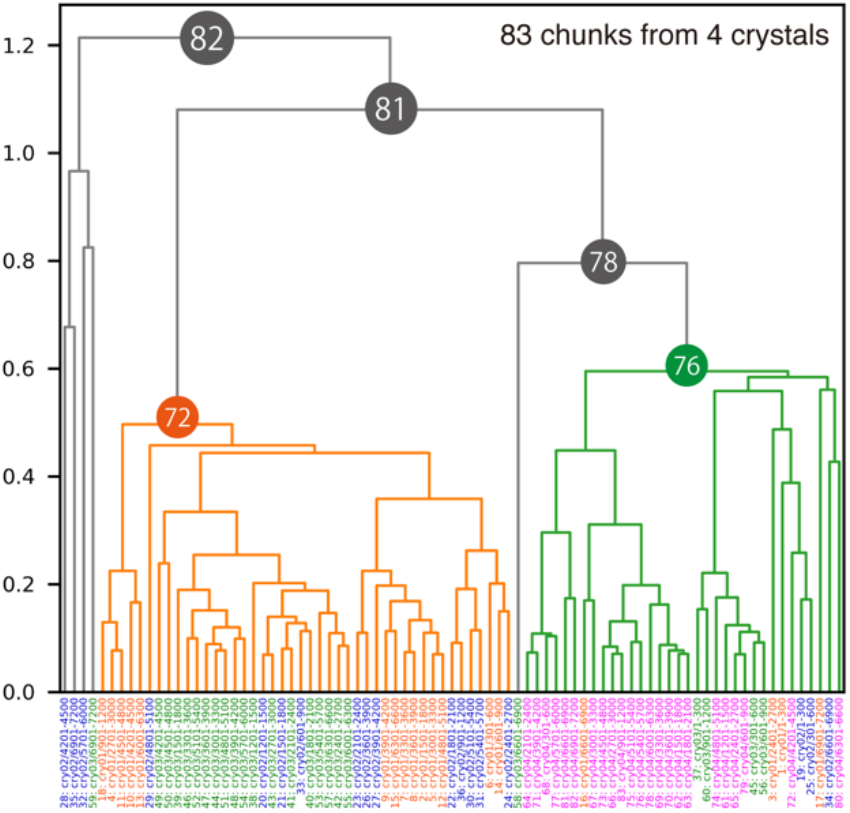
Dendrogram from intensity-based HCA for the Trn1-peptide complex. Data labels are shown at the bottom and colored by crystals. Clusters 72 and 76 were used for further structural analysis because these two clusters appeared to have different structures (polymorphs).

The electron density maps from these two nodes were significantly different (Fig. 7). There were two peptide-binding forms: one without any secondary structure (Form1) and the other with α-helix (Form2). These two peptide-binding forms were also observed when all chunks merged (Cluster 82). However, the *F*_o_-*F*_c_ map for each peptide-binding form appeared to be less clear than the clustering results (Clusters 72 and 76). Although both peptide-binding forms appeared in each node, occupancies seemed different. The *F*_o_-*F*_c_ map indicated that Form1 was dominant at Cluster 76 while Form2 was dominant at Cluster 72. The isomorphic threshold values range from 0.73 to 0.85, and looking at the dendrogram with these values, clusters 72 and 76 are candidates for structural polymorphs, consistent with the above results.

**Figure 7.**
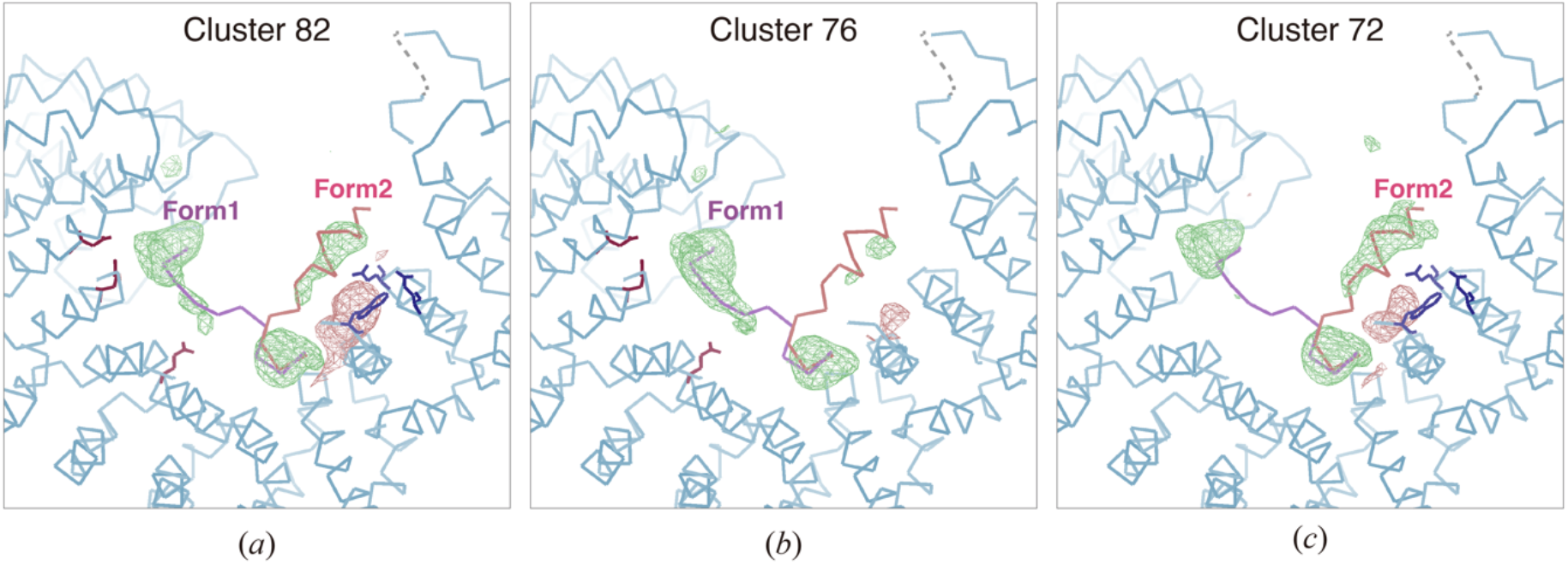
Peptide-omitted *F*_o_-*F*_c_ map from different nodes resulted from intensity-based HCA for the Trn1-peptide complex. *F*_o_-*F*_c_ maps calculated from merged data at (a) Cluster 82, (b) Cluster 76, and (c) Cluster 72 are depicted. The contour level for each figure is set to 3.0α. Binding peptide was omitted during the map calculation. Figures were generated by *Coot*.

The existence of two peptide binding forms was supported by the result of biochemical experiments. As illustrated in Fig. 7, some acidic amino acid residues were found around the *F*_o_-*F*_c_ density of the NLS peptide in both binding forms. Since NLS generally consists of basic amino acid residues, this is in line with the cargo recognition mechanism which is achieved by electrostatic interaction. Triple mutation for each peptide binding form (2 Glu and 1 Asp residues for Form1, and 2 Glu and 1 Trp residues for Form2) caused a decrease in peptide binding. Thus, both peptide binding forms are physiologically important for the function of Trn1.

Since for the Trn1-peptide complex, the variation of unit cell constants was more significant than that of trypsin (Fig. S14), we also applied unit cell-based HCA (Fig. S15). The largest LCV value for the Trn1-peptide complex was 4.03%, which was significantly higher than that for trypsin (0.59%). The obtained electron density maps seemed different (Fig. S16), as observed in the results from intensity-based HCA (Fig. 7). However, the clustering result was slightly different. Form1-dominant nodes appeared in the left branch (Clusters 73 and 78), while both forms were observed in the right branch (Clusters 77 and 79). The Form2 dominant node was not found in the result from unit cell-based HCA. Accordingly, a single-step intensity-based HCA seemed to classify the data better in the case even though the cell variation is significant.

Although the resolution was relatively low (lower than 3 Å), and the space group was not highly symmetric (*C2*) in the case of the Trn1-peptide complex, intensity-based HCA was adequate to identify two different peptide binding modes. The results imply that polymorph analysis can even be performed on datasets exhibiting relatively low resolution and low symmetry space groups.

Another interesting finding is that the dominant peptide binding form was different even though the crystals were obtained in the same crystallization drop. Furthermore, some crystals had both Form1-dominant and Form2-dominant chunks (*i.e.*, intra-crystal variation). Mapping the clustering results revealed that Form1-dominant chunks were mostly found at the tip of the crystals (Fig. S17). This result implies that polymorphs not only in different crystals (inter-crystal) but also in the same crystals (intra-crystal) are discernible by collecting diffraction data via the continuous helical scheme, significantly expanding the possibilities of polymorph analysis.

#### 3.2.2. Diffraction data classification and polymorph analysis on *Aa*HypD-C360S

Using a helical scheme, diffraction datasets of *Aa*HypD-C360S were acquired from six crystals applying 360° rotation per crystal. Each dataset was divided into 30° chunks, yielding 72 chunks from six crystals. The resolution for each chunk was approximately 1.6 Å. The number of reflections in each 30° chunk was about 100,000 reflections, overall completeness was about 45% and the space group was *P*2_1_2_1_2_1_. For the intensity-based HCA, reflections up to 2.79 Å were used to calculate the correlation between the datasets.

The values of the isomorphic threshold were about 1.9∼2.2 based on the dendrogram in Figure 8, and polymorph candidates were clusters 61, 63, and 64 in terms of this threshold. Since the Ward distance was very large compared to other samples in this paper, we decided to carefully perform structural analysis on the clusters for which complete datasets were available and compare the details of this sample. Among the data from these clusters, some polymorphs that have differences around the N-terminal and the [4Fe-4S] regions were identified by examining the corresponding electron density maps (Figs. 9 and S18). Significant differences were found in Clusters 50 and 52. At Cluster 50, the N-terminal region was unfolded (referred to as ‘unfolded’ conformation) with no secondary structure and the occupancy of the [4Fe-4S] cluster decreased with the disorder of the surrounding area (Fig 9a and 9b). Whereas, at Cluster 52, the N-terminal region was folded towards the protein side (referred to as ‘folded’ conformation), and the surrounding region around the [4Fe-4S] cluster was well-ordered (Fig. 9c and 9d). The electron density map in Cluster 51 (Fig. S18a and S18b) exhibited a similar trend in Cluster 50. However, the negative *F*_o_-*F*_c_ peak of the [4Fe-4S] cluster was decreased in Cluster 51. The electron density map in Cluster 42 was similar to that in Cluster 52 (Fig. S18c and S18d). However, the ‘folded’ N-terminal region was more evident in Cluster 52. Two different N-terminal conformations were highly related to the disorder around the [4Fe-4S] cluster. *B*-factor analysis clearly indicated that ‘unfolded’ conformation in N-terminal destabilized [4Fe-4S] surrounding region (Fig. 10). From data at Cluster 54, a significantly high *R*_free_ solution (*R*_free_ > 0.4) was obtained. At Cluster 47, complete data was not obtained because most chunks were rejected by outlier rejection in *KAMO*. In this example, the most significant structural differences (N-terminal and [4Fe-4S] region) appeared to be separated at the first branch of the dendrogram. During further separation, more slight differences (occupancies) seemed to be classified (Clusters 50 and 51, or Cluster 52 and 42).

**Figure 8.**
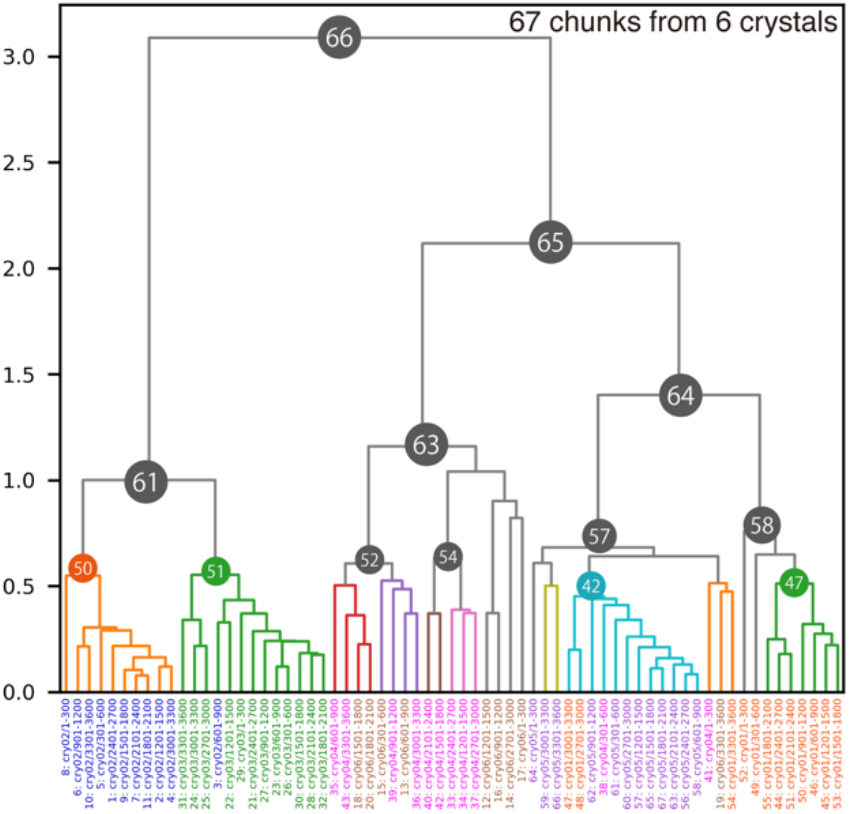
Dendrogram from intensity-based HCA on the *Aa*HypD-C360S variant. Data labels are shown at the bottom, colored by crystals. Due to the significant deviation among datasets, some clusters were not satisfied with enough completeness for further analysis. Accordingly, Clusters 52 and 54 were selected based on the results with the threshold of 0.8. Each data cluster consists of chunks from one or two crystals, indicating that inter-crystal differences are more prominent than intra-crystal differences in the *Aa*HypD-C360S.

**Figure 9.**
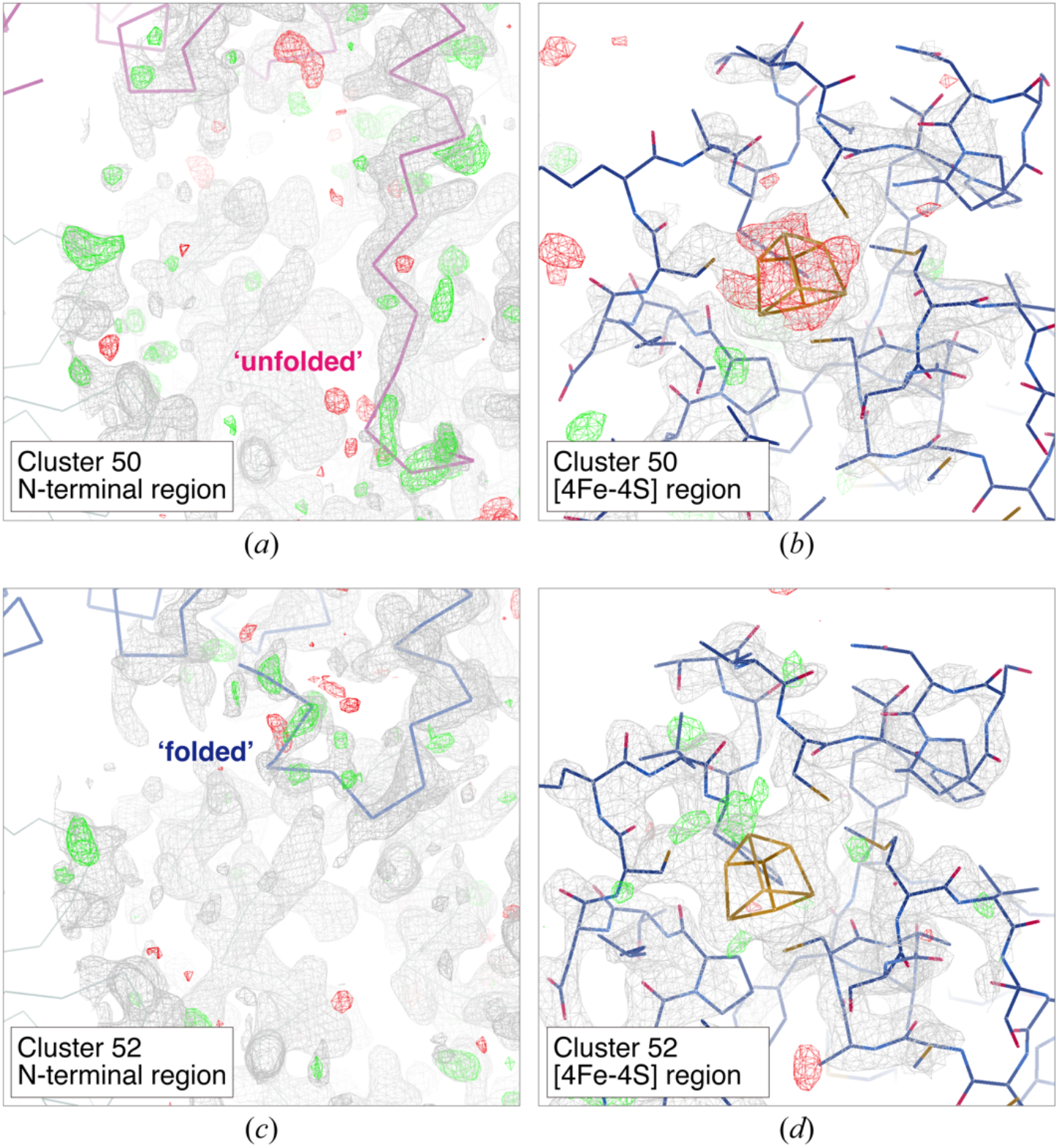
Electron density map around N-terminal region and [4Fe-4S] cluster obtained from merged data at different clusters: (a)-(b) Cluster 50 and (c)-(d) Cluster 52. The contour level for the 2*F*_o_-*F*_c_ map (gray mesh) is set to 1.00 except for the [4Fe-4S] region in Cluster 52, where that is set to 1.50. The contour level for the *F*_o_-*F*_c_ map is set to 3.00 (green mesh: positive, red mesh: negative) in all figures. Figures were generated by Coot. Only the main chain is depicted in the N-terminal region: ‘unfolded’ (purple), and ‘folded’ (blue). The variable N-terminal region (Ser7-Tyr12) was omitted and the occupancy for [4Fe-4S] cluster was set to 1.0 in map calculation. Figures were generated by *Coot*.

**Figure 10.**
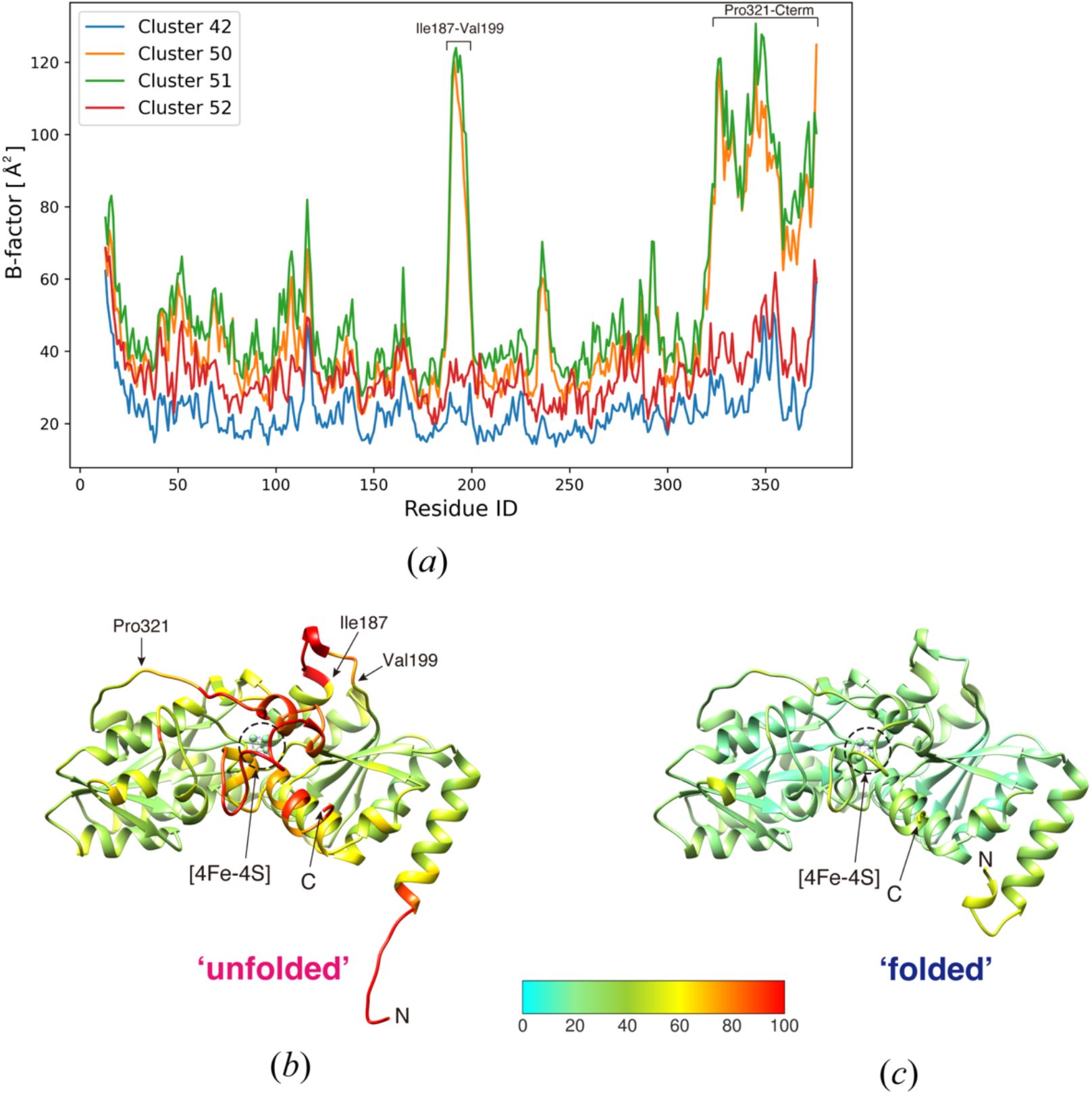
*B*-factor analysis on *Aa*HypD-C360S structures obtained from different clusters in intensity-based HCA. (a) Plot for *B*-factor of Cα atom in data from different clusters. (b)-(c) main-chain trace of *Aa*HypD-C360S with ‘unfolded’ N-terminal obtained from Cluster 50 (b) and with ‘folded’ N-terminal obtained from Cluster 52 (c). *B*-factor based coloring was applied using *Coot*. N-terminal and C-terminal are depicted in the figure as ‘N’ and ‘C’, respectively. The structure with ‘unfolded’ N-terminal showed a significantly high *B*-factor around the [4Fe-4S] cluster. Figures were generated by *UCSF Chimera* (Pettersen *et al*., 2004).

Consequently, clusters 61, 63, and 64, classified from our "isomorphic threshold," are the parent nodes of the data where the characteristic structures were found. Cluster 61 is the parent node for 50 and 51, cluster 63 for 52, and cluster 64 for 42 (Figure 8). Considering only locally for cluster 61, *W*_0_ is 1.0, corresponding to an isomorphic threshold of 0.6 to 0.7. This allows clusters 50 and 51 to be considered classifiable within that cluster. Our isomorphic thresholds proved to be valid also for this sample. Polymorphs were also found in crystals obtained under the same crystallization conditions. This result indicates that a protein may have several metastable conformations even under identical crystallization conditions. The results of this study indicate that when the absolute value of Ward distance on the dendrogram is large, as in the present sample, it is useful to try to analyze the structure of the subclusters of the nodes that diverged at the isomorphic threshold.

The median values of CC distributions for homologous data pairs in Clusters 50, 51, 52, and 42 were 0.976, 0.961, 0.935, and 0.966. The corresponding standard deviations were 0.019, 0.019, 0.020, and 0.019, respectively. These statistics are filtered by *d_CC_* <0.4 as well as other samples. As expected from the resultant dendrogram (Fig. 8), there are significant CC changes between the data pairs from Cluster 50 or 51 and Cluster 52 or 42. Even for the CC distribution for the data pairs between Cluster 50 and Cluster 51, where the smallest difference was found from the dendrogram, the mean value of the CC distribution was 0.946, and the corresponding standard deviation was 0.020.

### 3.3. Current limitations of intensity-based HCA and conceivable best practices for further applications

The results of the two representative samples demonstrated that intensity-based HCA with the proposed ‘isomorphic threshold’ can be a useful indicator to detect polymorphs in datasets obtained from multiple crystals. In addition, splitting the data collected by helical scheme into several chunks could be beneficial for analyzing structural polymorphs within the same crystals. Even for well-known proteins whose structures have already been determined, structural polymorphs have possibly been overlooked. The scope of this guideline is intended for cases where multiple crystals of 50 µm or larger are used to the extent that complete data can be collected from a single crystal with slight variation and cell constants. We focus on finding polymorphs by collecting large wedge datasets of 360 degrees or more through helical data collection from multiple large crystals and clustering them into chunk datasets of 30 degrees or more.

There is an explicit limitation for intensity-based HCA. A certain number of common reflections are required among the datasets for calculating CC value. As a default setting in *KAMO*, at least three common reflections are required between each dataset. In SWSX, diffraction data is collected for 5°–20° rotation from each crystal, which results in fewer common reflections. The number of common reflections also decreases due to the lower resolution or crystallographic symmetry. In this study, the chunk size was set at 30°, where the number of common reflections is expected to be adequate.

To investigate the limitation on chunk size (rotation range), the number of common reflections and rejected data in different chunk sizes were plotted using the high-resolution trypsin dataset from 0.5° to 30° (Fig. S19). The number of chunks excluded from the CC calculation was drastically increased below 3°. When the chunk size was set to 0.5°, almost all data were rejected (99.0%). The result is consistent with the previous report on intensity-based HCA applied to SSX (serial synchrotron crystallography) data (2°/crystal) using *ccCluster* (Santoni *et al*., 2017). Obviously, it is not realistic to apply the intensity-based HCA to single-frame data with extremely narrow or no oscillation data, such as collected by SFX (serial femtosecond crystallography) or SSROX (serial synchrotron rotation crystallography) approaches.

In addition, to investigate a sufficient number of common reflections for intensity-based HCA, the number of common reflections was evaluated in different chunk sizes for the datasets used in this study. Since the rejected data increased below 3° in the case of trypsin, log (Number of common reflections) ≧ 2.5 seems promising. Fraction of common reflections in total reflections are highly dependent on crystallographic symmetry (Fig. S20). If the resolution is relatively low or crystallographic symmetry is not high, a large molecular weight could cover the number of reflections, as exhibited in the example of Trn1-peptide complex. If possible, partial data collection by the helical scheme or larger rotation data will help increase the common reflections for polymorph analysis with intensity-based HCA.

Unit cell-based HCA can still be useful when intensity-based HCA is not available. Even for single-frame data, such as SFX data, unit cell constants are available. Therefore, unit cell-based HCA can be applied to any given diffraction dataset. In actual SFX data, datasets with different unit cell properties are sometimes mixed in the entire datasets (Nomura *et al*., 2021). Considering a general case, unit cell-based HCA should first be applied to filter the dataset with different cell parameters. If the variation of unit cell constants is relatively small, say the largest LCV value is less than 1%, it is likely that unit cell-based HCA will not yield good classification results. However, polymorphs could be found from the result of unit cell-based HCA. In a recent study (Soares *et al*., 2022), the distance metric for unit cell-based HCA has been improved. Furthermore, outlier rejection during the merging step as implemented in *KAMO* could be useful to reduce contaminated dataset.

Although we classified polymorphs by a single-step intensity-based clustering, cell parameters should be considered for more accurate classification in general cases. From the result of unit cell-based HCA for trypsin datasets, chunks from the same data were separated into different branches (Figs. 1 and 3). If intensity-based HCA is further applied to the clusters obtained from unit cell-based HCA, the number of datasets remaining at the final stage of data merging may decrease. Thus, single-step clustering is preferable when the total number of datasets is small. HCA with 2-dimensional parameters, including both cell parameters and intensity correlation, is under consideration for more effective single-step clustering.

## 4. Conclusions and outlook

Based on the investigations on the test cases, we propose that the "isomorphic threshold" for classification by intensity-based HCA of several hundred 30° chunk data is the Ward distance in the top row of the dendrogram multiplied by 0.6-0.7. In the representative samples, polymorphs were successfully detected by intensity-based HCA with our suggested threshold. The scope of this guideline basically includes the cases where multiple crystals larger than 50 µm are used and complete data can be collected from each single crystal with slight variation in unit cell constants. We focus on finding polymorphs by collecting large wedge datasets of 360 degrees or more through helical data collection from multiple large crystals and clustering them into chunk datasets of 30 degrees or more.

The results of the study, including standard and representative samples, show that the single-step intensity-based HCA with the proposed "isomorphic threshold" is effective for detecting polymorphs using multiple datasets. Although helical data collection and HCA have been used in MX experiments and analyses, we demonstrated several advantages consistent with current HDRMX trends. The helical data scheme is unique in that it can be applied to polymorph analysis within crystals by dividing the complete dataset into chunks, as the data is collected while translating the X-ray exposure position. Indeed, our results suggest the possibility of the presence of polymorphs within crystals. The continuous helical data collection adopted in the automated data acquisition system, ZOO, at SPring-8 collects data from each crystal volume with a uniform dose, which may reduce structural inhomogeneities due to radiation damage compared to single-point rotation. Even when multiple crystals are used, the isomorphism among crystals due to radiation damage can be suppressed to the same extent, allowing discussion of only the structural differences that exist in the crystals. In addition, highly efficient automated data collection increases the number of datasets that can be measured in a given time, allowing the analysis of many polymorphs with higher resolution.

Our findings could be widely applied to other protein samples. Even though the resolution was relatively low (approximately 4 Å), physiologically meaningful polymorphs can be identified as exhibited in the example of Trn1-peptide complex. The representative examples also showed that protein molecules may exhibit different structures within the same crystal. In addition, polymorphs can be found from multiple crystals obtained under the same crystallization conditions in both practical cases. Determination of the ‘isomorphic’ threshold enables polymorph detection, even if the existence of polymorphs is not expected or detected during the sample preparation. It will also facilitate broader applications by automating the polymorph analysis with our proposed ‘isomorphic’ threshold.

The suggested polymorph analysis could help obtain various structural snapshots during the protein function process for elucidating the molecular mechanism. The determination of multiple structural snapshots will also contribute to more accurate structure prediction, as realized by *AlphaFold2* (Jumper *et al*., 2021) or *RoseTTAFold* (Baek *et al*., 2021). However, structural information on time series or reaction pathways among the identified polymorphs is not available by our approach. It may be complemented by molecular dynamics (MD) simulations. For instance, the free energy landscape analysis will help understand the dynamic structural mechanism during protein function (Oide *et al*., 2020). Polymorph analysis will be further enhanced along with the techniques for inducting structural change, such as ligand mixing. Although time-resolved crystallography is a powerful tool for analyzing dynamics, the proposed polymorph analysis will help compensate for numerous difficulties in controlling such reactions. Expanding the intensity-based HCA for single-frame data (from SFX or SSROX) should also be developed in the near future.

## 5. Availability

The here presented analysis can be performed anywhere by installing the program *KAMO*, which is available on GitHub (https://github.com/keitaroyam/yamtbx). The raw diffraction data used in this study are available from Zenodo.

## Supporting information

Supplemental sections/figures/tables.

## Acknowledgements

We thank Keitaro Yamashita (MRC, LBL) for the helpful discussions, and Mr. Kaede Nakayama (University of Hyogo) for technical assistance on simulations of HCA. We also thank the BL45XU beamline staff for assisting with X-ray crystallographic data collection. We sincerely thank Chai Gopalasingam and Christoph Gerle for critical reading of the manuscript and helpful suggestions. This research was supported by a Grant-in-Aid from the Japanese Ministry of Education, Culture, Sports, Science, and Technology (21K06031) (STF), a Grant-in-Aid for Scientific Research on Innovative Areas Grant numbers JP19H05762 (SA), 19H05783(MY), the Japan Society for the Promotions of Science KAKENHI Grant Numbers, JP20K06517 (NM), AMED-CREST under Grant Number 22gm1410010s0202 (STF), and the Platform Project for Supporting Drug Discovery and Life Science Research (Basis for Supporting Innovative Drug Discovery and Life Science Research (BINDS)) from AMED under Grant Number JP21am0101072 (support number 1587) (STF) and JP20am0101070 (MY).

