## Supplemental sections/figures/tables. for "Elucidating polymorphs of crystal structures with intensity-based hierarchical clustering analysis on multiple diffraction datasets"

### Elucidating structural polymorphs in multiple diffraction datasets using hierarchical clustering

#### S1. Investigations on the linkage methods for intensity-based clustering

The appearance of the dendrogram obtained from the result of hierarchical clustering can be completely different by the choice of the linkage method. Seven linkage methods are available in *scipy* module, namely, single, complete, average, weighted, centroid, median, and Ward methods. These seven linkage methods were applied to the distance matrix generated by *KAMO* during the intensity-based HCA on *in silico* mixed datasets including apo- and benzamidine-bound trypsin (Fig. S3). Dendrograms were colored without setting color threshold except for that with 'Ward' linkage. Although clusters with homogeneous dataset were appeared to be obtained with each linkage method, only the result with "Ward" linkage exhibited clear separation of two different datasets.

#### S2. Details in HCA simulation

We attempted to establish how to estimate the isomorphic threshold by HCA simulation using CC model. Assuming two structural polymorphs are mixed, three CC distributions should be considered (Fig S4). Two of these are homogeneous data pairs ($CC_{A\text{-}A}$ and $CC_{B\text{-}B}$) and the other is for heterogeneous ($CC_{A\text{-}B}$). In our simulation, the CC distributions were modelled by log-normal distribution by fitting to the observed datasets. We used a Python script, utilizing the *scipy* module. Specifically, we utilized the *scipy.stats.lognorm* with the following triplet parameters, (*s*, *loc*, *scale*) (See also a documentation page of *scipy.stats.lognorm*). The parameters for a CC distribution were estimated by the least-squares fitting of *lognorm.pdf(1-CC, s, loc, scale)* to corresponding observed CC distribution. Observed CC values above 0.8 were used to cover both the major peak and the tail. In the simulation, the CC values for CC matrix were extracted randomly from the corresponding model treated as a probability density function. For example, the CC value between A and B, the $CC_{A\text{-}B}$ model was chosen, and CC was randomly extracted to create the distance matrix (Fig. S3).

The two simulations were performed based on the CC models of trypsin. Firstly, assuming 50 datasets for each of the two structural polymorphs, HCA was performed with the parameters (*s*, *loc*, *scale*) estimated from observed datasets to evaluate whether the simulation is valid. After 100 repetitions, the mean and standard deviation of the Ward distance for classifying the two polymorphisms were calculated ($W_1$ in Fig S4).

Secondly, using the CC model, we tried to find a good R in equation 1. If the structures of apo- and benzamidine bound trypsin become very similar, the distributions of CC should overlap, making classification difficult. In the CC model, *loc* parameter represents the location of the CC distribution. In this simulation, the distributions of $CC_{apo-benz}$ and $CC_{benz-benz}$ were hypothetically brought closer by gradually moving the *loc* parameter of $CC_{apo-benz}$ closer to that of $CC_{benz-benz}$. Here, we divided the difference between the original $CC_{apo-benz}$ and $CC_{benz-benz}$ *loc* parameters by ten and repeated the HCA simulation for ten steps, moving the $CC_{apo-benz}$ *loc* parameters closer to $CC_{benz-benz}$ by one step. We performed HCA simulation assuming that there were an equal number of apo- and benz-trypsin datasets and evaluated whether they could be neatly classified into two on the dendrogram. For the evaluation, we defined a score that quantifies whether apo- and benz-trypsin can be classified into two clusters on the dendrogram.

Two types of indicators are considered: *balance score* and *purity score*. The balance score, assuming that the HCA simulation can divide datasets into two clusters and the number of data in each cluster is $n_1$ and $n_2$.

$$\textit{balance score} = 1 - |n_1 - n_2| / (n_1 + n_2)$$

where n1 and n2 are the number of datasets in classified two clusters. Since we assume an equal number of two structures in this simulation, this score should be 1.0 if the HCA classification is successful.

Purity is the percentage of A or B structure in each cluster, a number between 0 and 1, where 1 indicates 100% purity. We compared the purity of A and B in cluster 1 and set the larger of the two as the score, $s_1$, for cluster 1. Similarly, the greater of the purity of A and the purity of B in cluster 2 was set as the purity of cluster 2, $s_2$. The *purity score* was defined as the smaller of $s_1$ and $s_2$.

Finally, the total score was defined as the sum of the balance score and the purity score as follows,

$$\textit{total score} = 0.5 * \textit{balance score} + 0.5 * \textit{purity score}$$

The "$W_0$" is the Ward distance of the top cluster, and $W_1$ is the larger Ward distance of the two clusters classified in the dendrogram. The value of "$W_1$" eventually equals to the isomorphic threshold (equation1 and Figure S3). By plotting the resulting scores against R, we selected a valid R to estimate the "isomorphic threshold." This simulation was performed for 100, 200, 300, 500, and 1000 data sets.

**S3. Preparation of Transportin-1 in complex with a nuclear localization signal peptide**

The Transportin-1 (Trn1) was produced using an *Escherichia coli* (*E. coli*) expression system. The pGEX6p1-hTrn1 plasmids were gifts from Dr. M. Sato. The DNA fragments of Trn1Δloop mutant in which a part of the H8 loop (Trn1 344–375) was replaced with the GGSGGSG linker was subcloned into pGEX6p-1. *E. coli* BL21 (DE3) RIPL cells were transformed with the vector and were cultured at 37˚C in LB medium to a suitable cell density ($OD_{600}$ approximately 0.6–0.7), and then induced protein expression with IPTG (final concentration 0.1 m*M*) overnight at 18˚C. Bacteria cells were collected by centrifugation at 4000 rpm for 20 min at 20°C, suspended in a sonication buffer [110 m*M* potassium acetate, 20 m*M* HEPES-KOH pH 7.3, 10 m*M* dithiothreitol (DTT), 1 m*M* EDTA, 1 m*M* phenylmethylsulfonyl fluoride (PMSF), 20% (*v/v*) glycerol], sonicated, and centrifuged at 15,000 × *g* for 20 min at 4°C. The supernatant of the bacteria was loaded onto Glutathione Sepharose 4B (Cytiva). After washing with a wash buffer [110 m*M* potassium acetate, 20 m*M* HEPES-KOH pH 7.3, 10 m*M* DTT, 1 m*M* EDTA, 20% (*v/v*) glycerol] the GST-tagged Trn1 proteins were eluted with an elution buffer [110 m*M* potassium acetate, 20 m*M* HEPES-KOH pH 7.3, 10 m*M* DTT, 1 m*M* EDTA, 20% (*v/v*) glycerol, 50 m*M* reduced glutathione]. The eluate was incubated with 20–25 mg $ml^{-1}$ of PreScission Protease (Cytiva) overnight at 4°C to cleave the GST-tag. The sample was then loaded onto a 5 ml HiTrap Q HP Column (GE Healthcare), and Trn1 was eluted by applying a gradient of Buffer A [110 m*M* potassium acetate, 20 m*M* HEPES-KOH pH 7.3, 10 m*M* DTT, 1 m*M* EDTA, 10% (*v/v*) glycerol] and Buffer B [1 *M* potassium acetate, 20 m*M* HEPES-KOH pH 7.3, 10 m*M* DTT, 1 m*M* EDTA, 10% (*v/v*) glycerol]. The eluted protein solution was concentrated with Amicon Ultra-15 (Merk Millipore) and purified by gel filtration column chromatography using a Superdex 200 pg (column volume = 24 ml, Cytiva) with gel filtration buffer (110 m*M* potassium acetate, 20 m*M* HEPES-KOH pH 7.3, 10 m*M* DTT).

An NLS peptide of Trn1 was purchased from Eurofins and dissolved in a gel filtration buffer. By mixing Trn1Δloop and the peptide solution, Trn1-peptide complex solution was prepared (the final concentration of Trn1Δloop and NLS peptide were 5 mg $ml^{-1}$ and 5 m*M*, respectively). Then, 1 µl of the Trn1- peptide complex solution and 1 µl of reservoir solution (0.5 *M* NaK phosphate pH 5.0) were mixed at room temperature, and crystallization was performed using the sitting-drop vapor diffusion method at 10˚C. The obtained crystals were cryoprotected by 30% (*w/v*) glycerol-containing reservoir solution prior to data collection.

**S4. Data statistics for merged data obtained at cluster nodes**

Crystallographic statistics values may help interpret the existence of polymorphs in the obtained datasets. Especially, some statistic values, *e.g.* <I/σI>, $R_{\mathrm{meas}}$, $CC_{1/2}$, obtained from merged datasets at each cluster from intensity-based HCA may be informative. These statistics are summarized in Table S4-S7 for four datasets (two kinds of *in silico* mixed trypsin datasets, Trn1-peptide complex, and *Aa*HypD-C360S used in this study), respectively. Resolution cut-off for each merged data was estimated using *kamo.decide_resolution_cutoff* with the $CC_{1/2}$ of outer resolution shell (approximately $CC_{1/2} \sim 0.50$). All the statistics values were obtained from output of *XSCALE* for finally obtained merged data after outlier rejection process of *KAMO*. Number of chunks (30° each) is obtained from input files for *XSCALE*, however, some frames may be rejected.

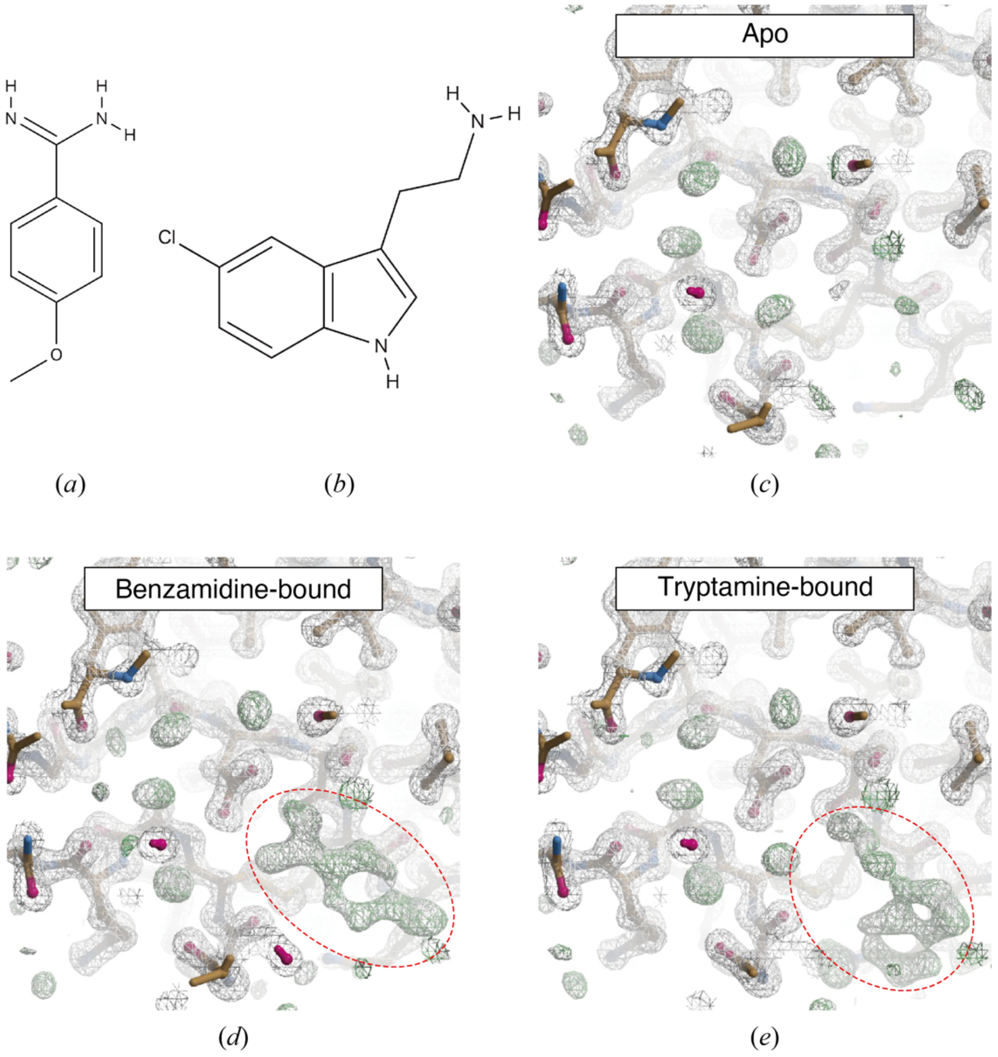

**Figure S1** $2F_o$-$F_c$ and $F_o$-$F_c$ map around the inhibitor binding site of trypsin from a large-wedge dataset without splitting into chunks. Here the results from only a single crystal are shown as a representative of each trypsin dataset. Structural formulas of inhibitors used in this study are shown in (a) 4-methoxybenzamidine (referred to as 'benzamidine') and (b) 5-chlorotryptamine (referred to as 'tryptamine'). $2F_o$-$F_c$ (gray mesh) and $F_o$-$F_c$ (green) density maps obtained from (c) apo-trypsin, (d) benzamidine-bound, and (e) tryptamine-bound trypsin datasets are depicted. Inhibitors were omitted in map calculation. The contour level for each $2F_o$-$F_c$ and $F_o$-$F_c$ map is 1.0σ and 3.0σ, respectively. Structural formulas for inhibitors were depicted by *Molview* (https://molview.org). $2F_o$-$F_c$ and $F_o$-$F_c$ maps were generated by *Coot*.

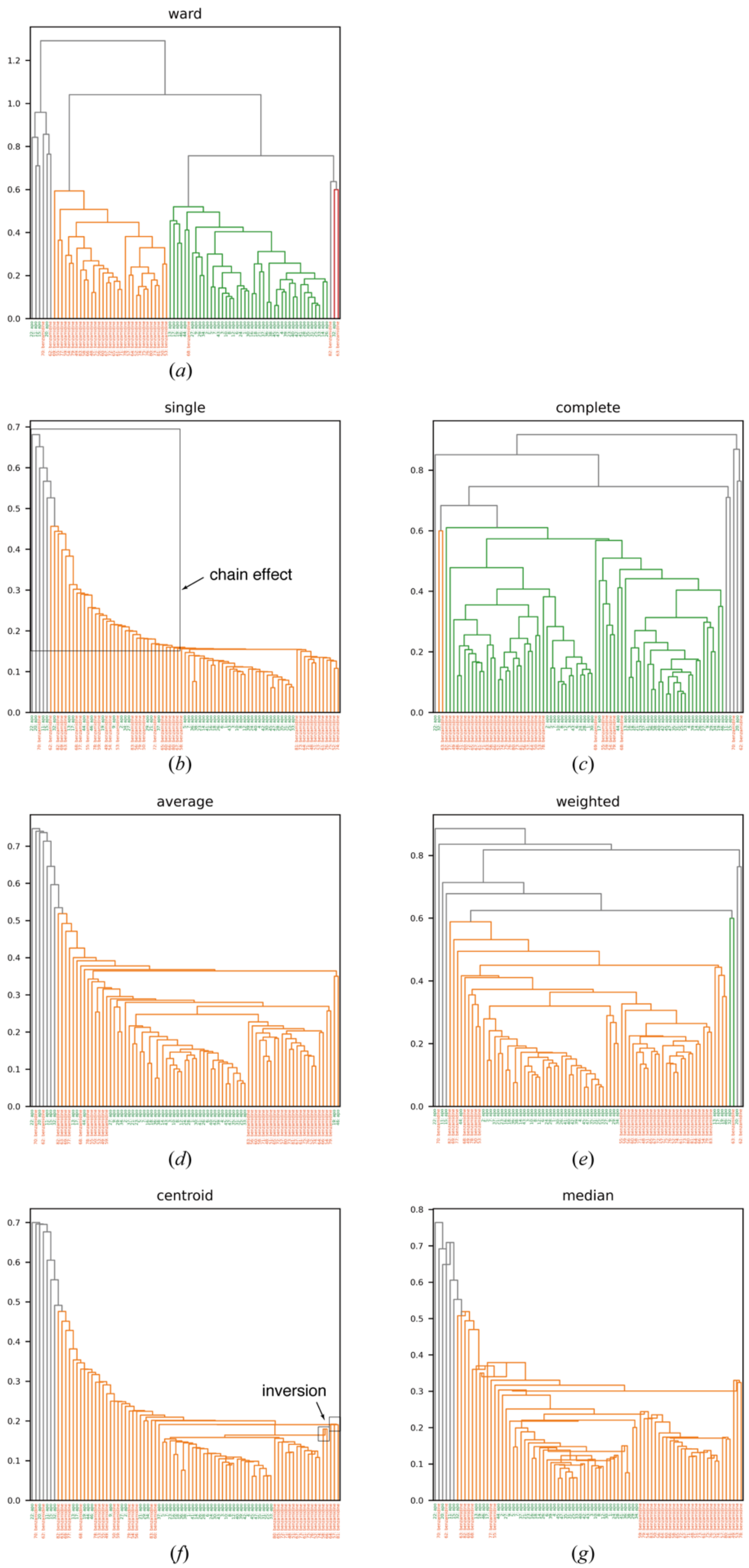


**Figure S2** Dendrogram obtained with different linkage method. The distance matrix was obtained from the intensity-based HCA on *in silico* mixed dataset including apo and benzamidine-bound trypsin. Seven linkage method, namely, “Ward”, “single”, “complete”, “average”, “weighted”, “centroid”, and “median”, available in *scipy* module was applied. Data label at each leaf is colored by green (apo-trypsin) and orange (benzamidine-bound trypsin). The color threshold for each dendrogram were not set (0.7 × maximum distance in default) except for the data with “Ward” linkage (used 0.6 as well as Fig. 2).

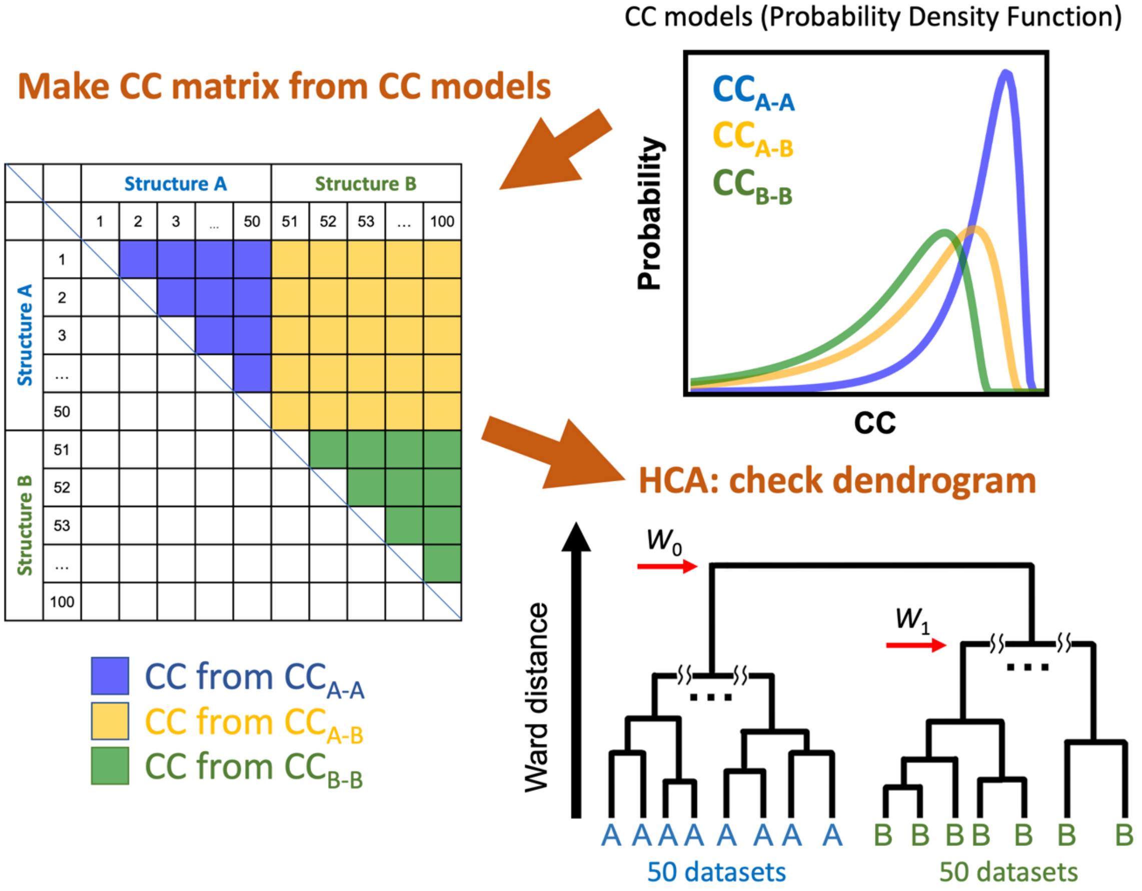


**Figure S3** Schematic diagram of our HCA simulation. We assume two mixed data sets, Structure A and Structure B, each with 50 datasets in this figure. We first fit a log-normal function to the observations to obtain probability density functions for each of the three CC distributions ($CC_{A\text{-}A}$, $CC_{B\text{-}B}$, and $CC_{A\text{-}B}$). CC values were randomly extracted from the models, and a CC matrix was created to compute the distance matrix. Each colored box in the table contains the extracted CCs and the color is the same as the color of the model used for extraction. HCA was performed from the distance matrix to see if the dendrogram classified the A and B structures. The isomorphic threshold is calculated as $W_1/W_0$ by using $W_0$ and $W_1$. Each value is Ward distance indicated by the red arrow.

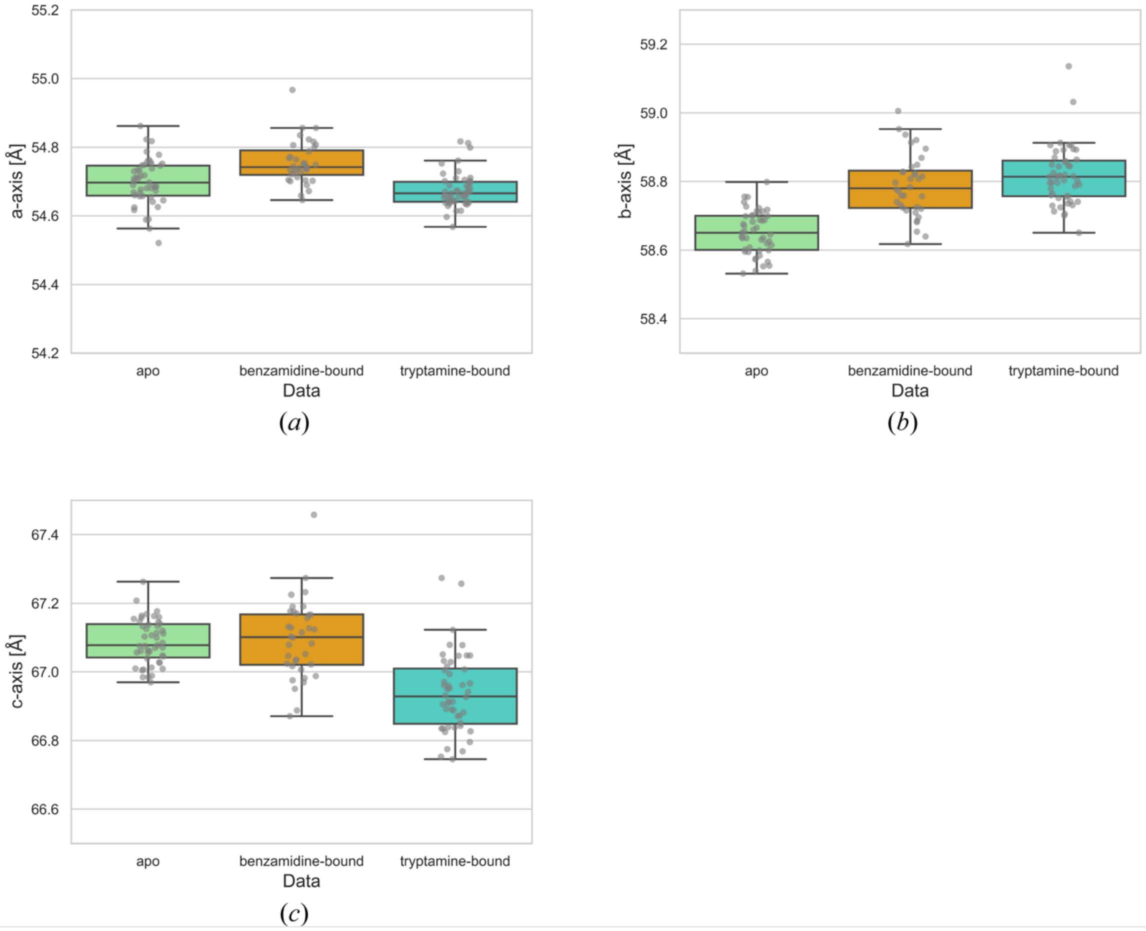


**Figure S4** Comparison of unit cell constants among apo-trypsin and inhibitor-bound trypsin datasets. The distribution of each cell parameters was illustrated with combination of box plot and swarm plot. The vertical axis of each plot was depicted with the same scale. Although all datasets were slightly different at each unit cell axis, the distribution of each axis value was largely overlapped.

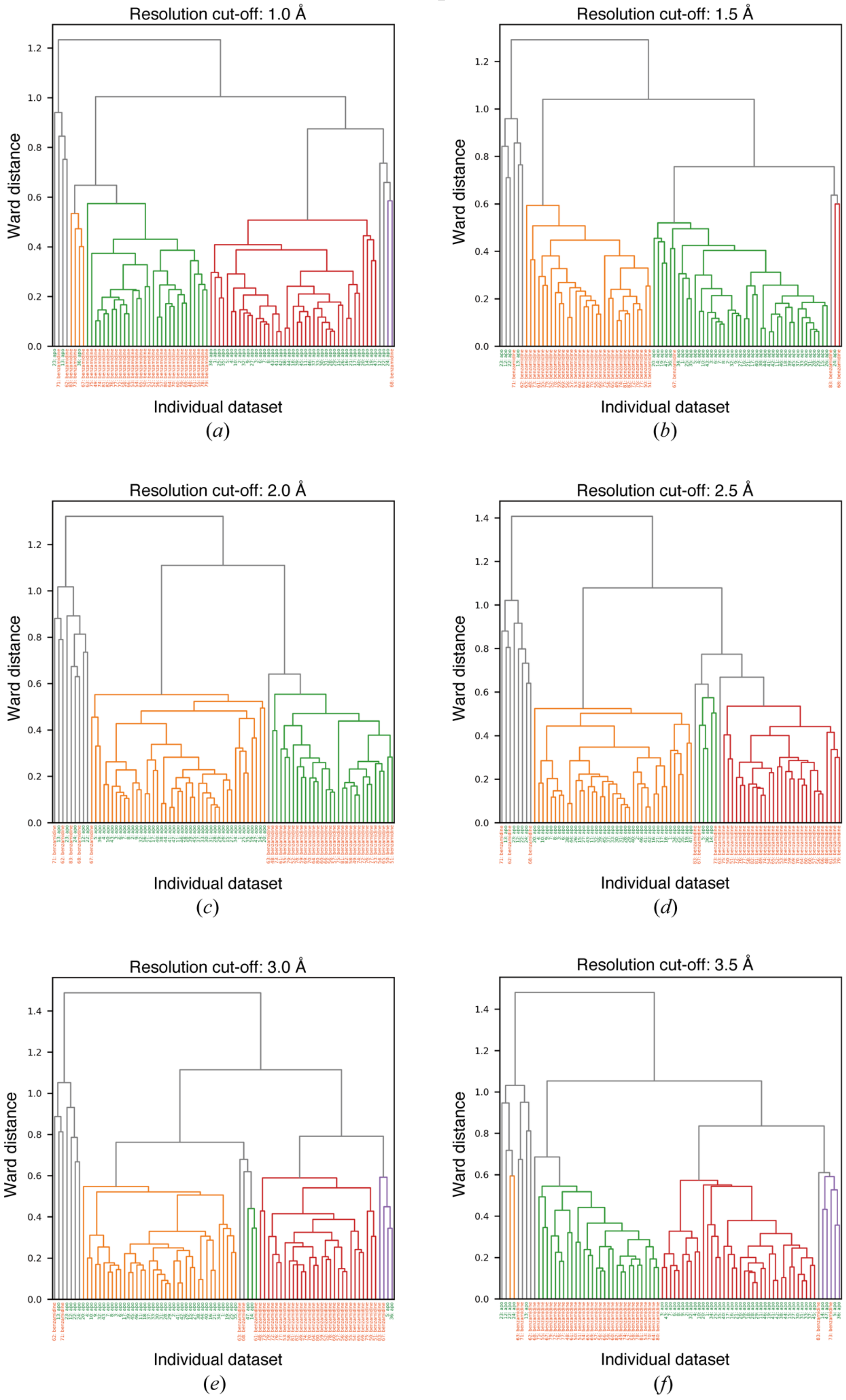

Resolution cut-off: 1.0 Å
Ward distance
Individual dataset
(a)
Resolution cut-off: 1.5 Å
Ward distance
Individual dataset
(b)
Resolution cut-off: 2.0 Å
Ward distance
Individual dataset
(c)
Resolution cut-off: 2.5 Å
Ward distance
Individual dataset
(d)
Resolution cut-off: 3.0 Å
Ward distance
Individual dataset
(e)
Resolution cut-off: 3.5 Å
Ward distance
Individual dataset
(f)

**Figure S5** Result of intensity-based HCA on in silico mixed datasets containing apo- and benzamidine-bound trypsin with different resolution cut-off. Leaf label was colored in green (apo) and orange (benzamidine-bound). Cut-off resolution for each panel is (a) 1.0 Å, (b) 1.5 Å, (c) 2.0 Å, (d) 2.5 Å, (e) 3.0 Å, (f) 3.5 Å, respectively.

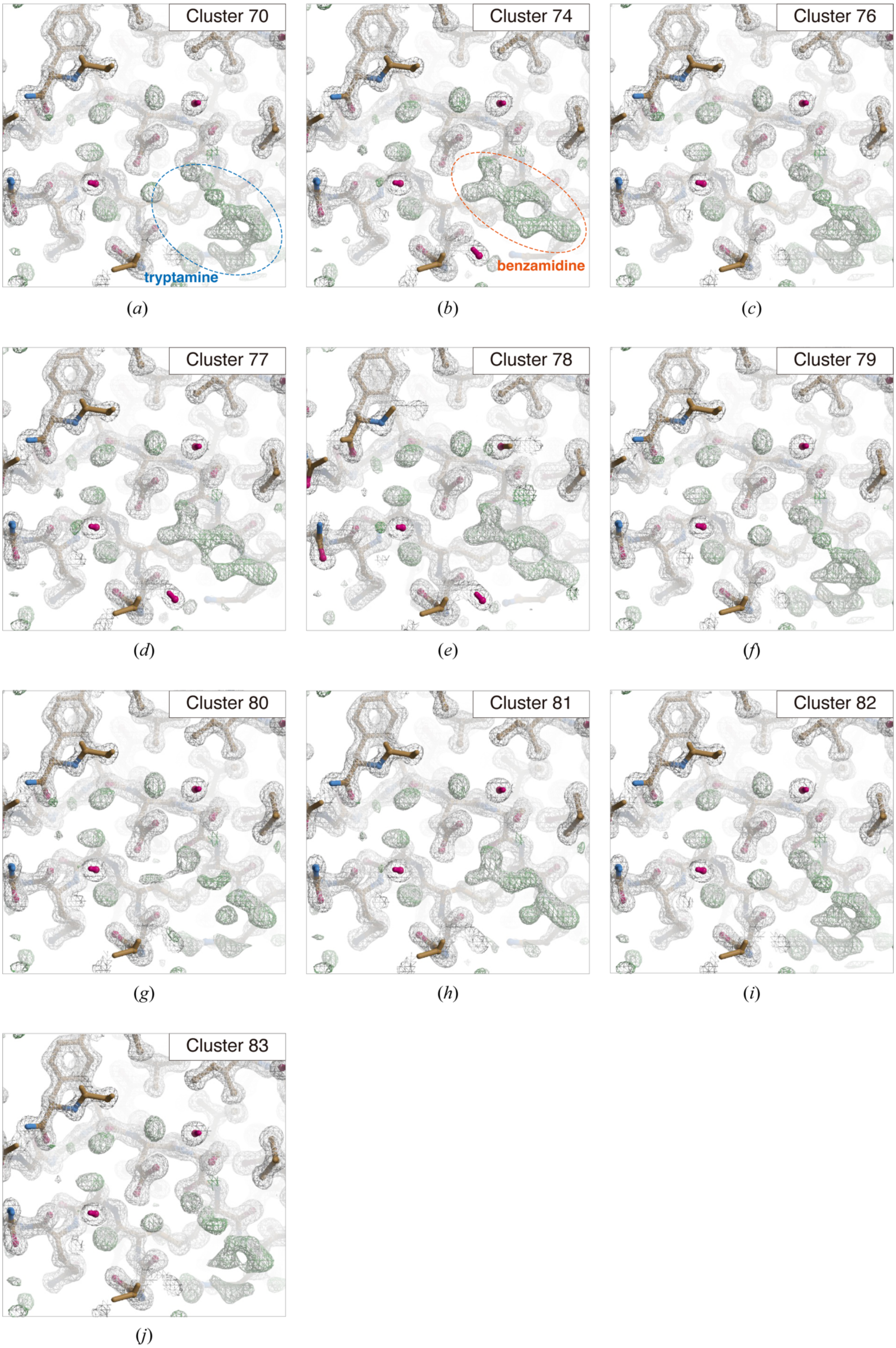

Cluster 70
tryptamine
(a)
Cluster 74
benzamidine
(b)
Cluster 76
(c)
Cluster 77
(d)
Cluster 78
(e)
Cluster 79
(f)
Cluster 80
(g)
Cluster 81
(h)
Cluster 82
(i)
Cluster 83
(j)

**Figure S6** Result of unit cell-based HCA on *in silico* mixed datasets containing benzamidine-bound and tryptamine-bound trypsin datasets represented by the electron density maps. Electron density maps calculated from merged data at different nodes in the dendrogram are illustrated. The contour level of $2F_o$-$F_c$ map (gray mesh) and $F_o$-$F_c$ map (green mesh) are 1.0σ and 3.0σ, respectively. Figures were generated by *Coot* exploited in the *NABE* pipeline.

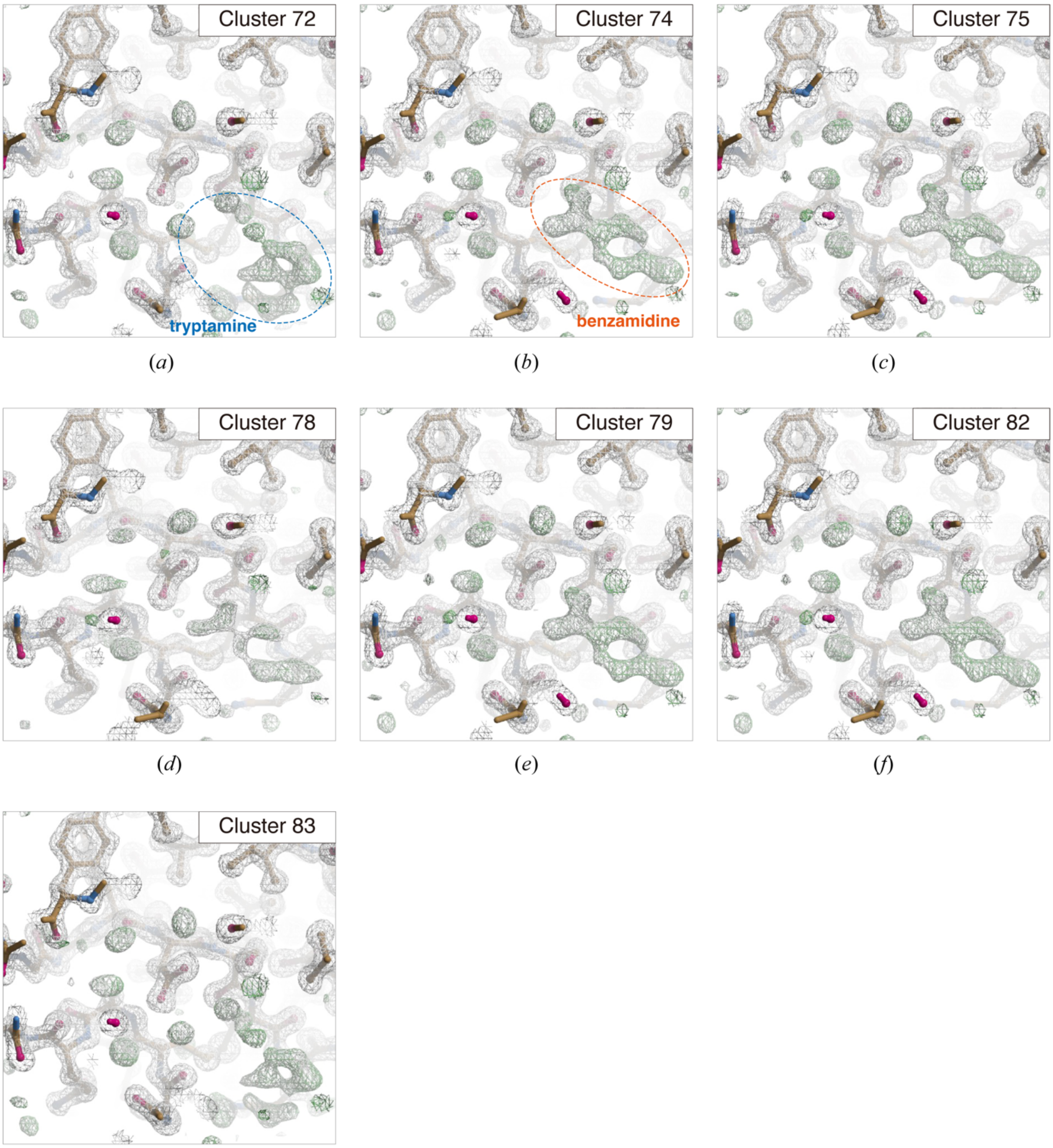


(*a*) (*b*) (*c*)

(*d*) (*e*) (*f*)

(*g*)

**Figure S7** Result of intensity-based HCA on *in silico* mixed datasets containing benzamidine-bound and tryptamine-bound trypsin datasets represented by the electron density maps. Electron density maps calculated from merged data at different nodes in the dendrogram are illustrated. The contour level of $2F_o$-$F_c$ map (gray mesh) and $F_o$-$F_c$ map (green mesh) are 1.0σ and 3.0σ, respectively. Figures were generated by *Coot* exploited in the *NABE* pipeline.

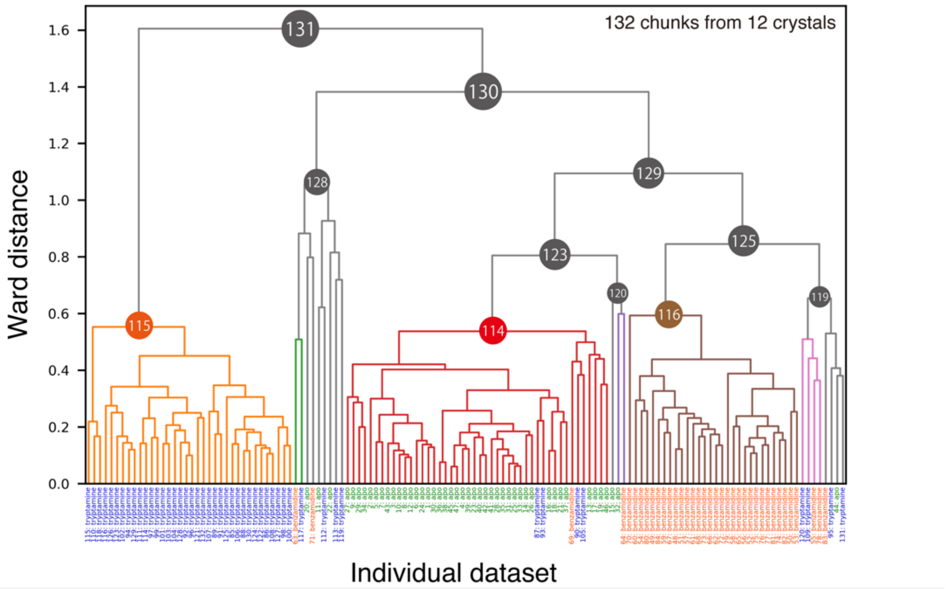


**Figure S8** Results of the intensity-based HCA on *in silico* mixed dataset containing apo-, benzamidine-bound, and tryptamine-bound trypsin. Cluster 115, 114 and 116 are obtained as isomorphic clusters within the suggested 'isomorphic threshold'.

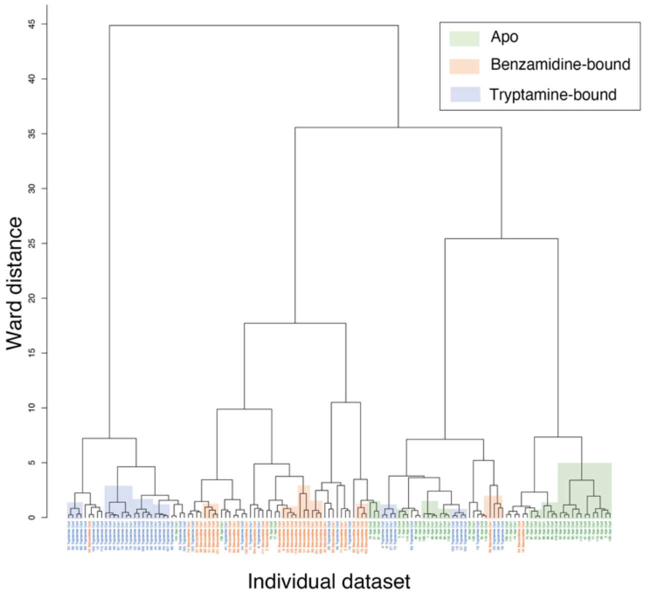

**Figure S9** Dendrogram from unit cell-based HCA on *in silico* mixed datasets consisting of apo-, benzamidine-bound, and tryptamine-bound trypsin. The color label for each dataset is set to green (apo-trypsin), orange (benzamidine-bound trypsin), and blue (tryptamine-bound trypsin). Cluster in the dendrogram is colored by the same color for the leaf label, when more than three chunks of same data make cluster.

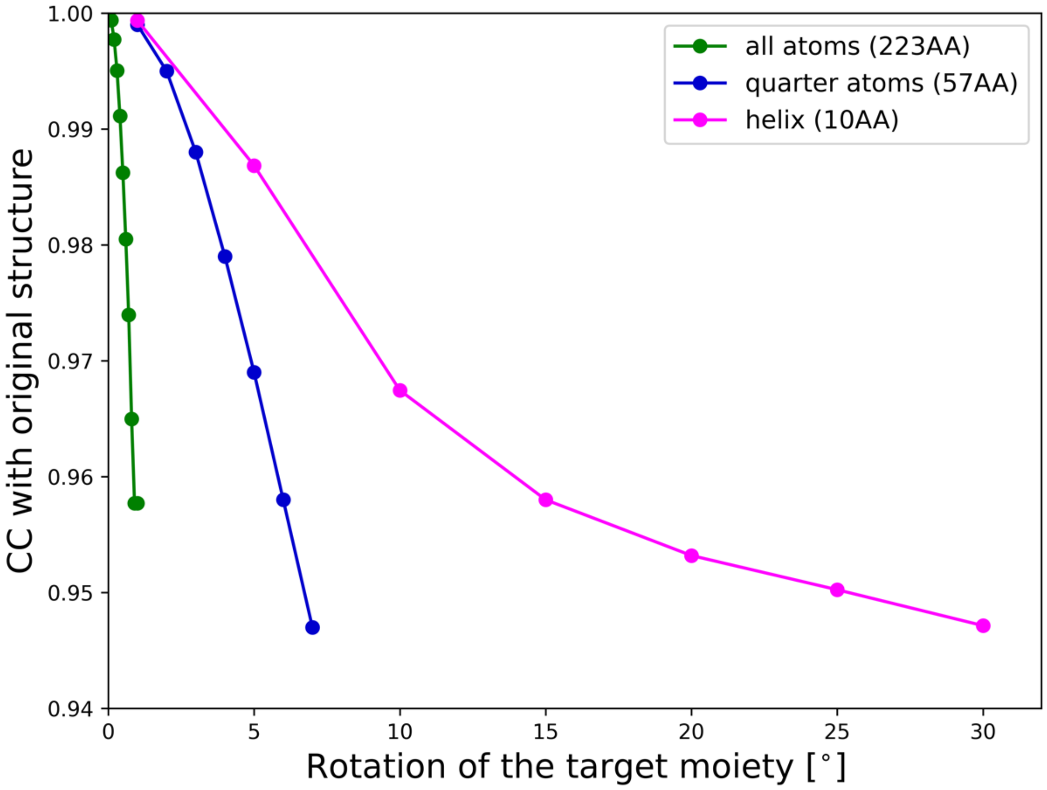


**Figure S10** Change of CC by rotation of a certain moiety of trypsin molecule. CC between the original and rotated trypsin molecule is plotted. The plot is colored by the rotated moiety; green: whole trypsin molecule (223 amino acid (AA) residues), blue: quarter of the molecule (57 AA), and magenta: a terminal helix (10 AA length).

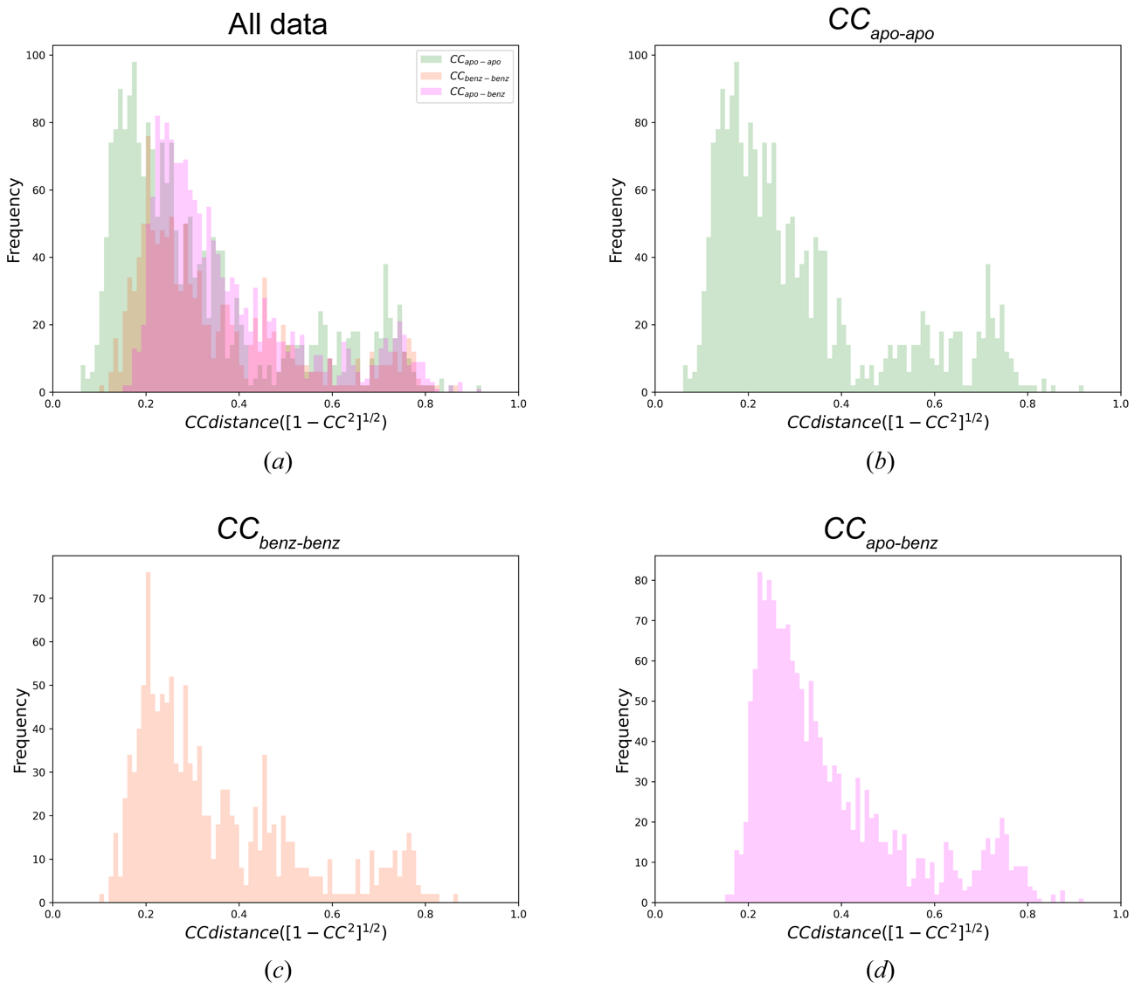


**Figure S11** Histogram of $d_{CC}$ (CC distance: $[1 - CC^2]^{1/2}$) calculated between homogeneous and heterogeneous chunks of trypsin. (a) All histogram are drawn in the same plot. (b) $d_{CC}$ obtained between two apo chunks $CC_{apo\text{-}apo}$, (c) $d_{CC}$ obtained between two benzamidine-bound chunks $CC_{benz\text{-}benz}$, (d) $d_{CC}$ obtained for heterogeneous combination of apo and benzamidine-bound chunk $CC_{apo\text{-}benz}$

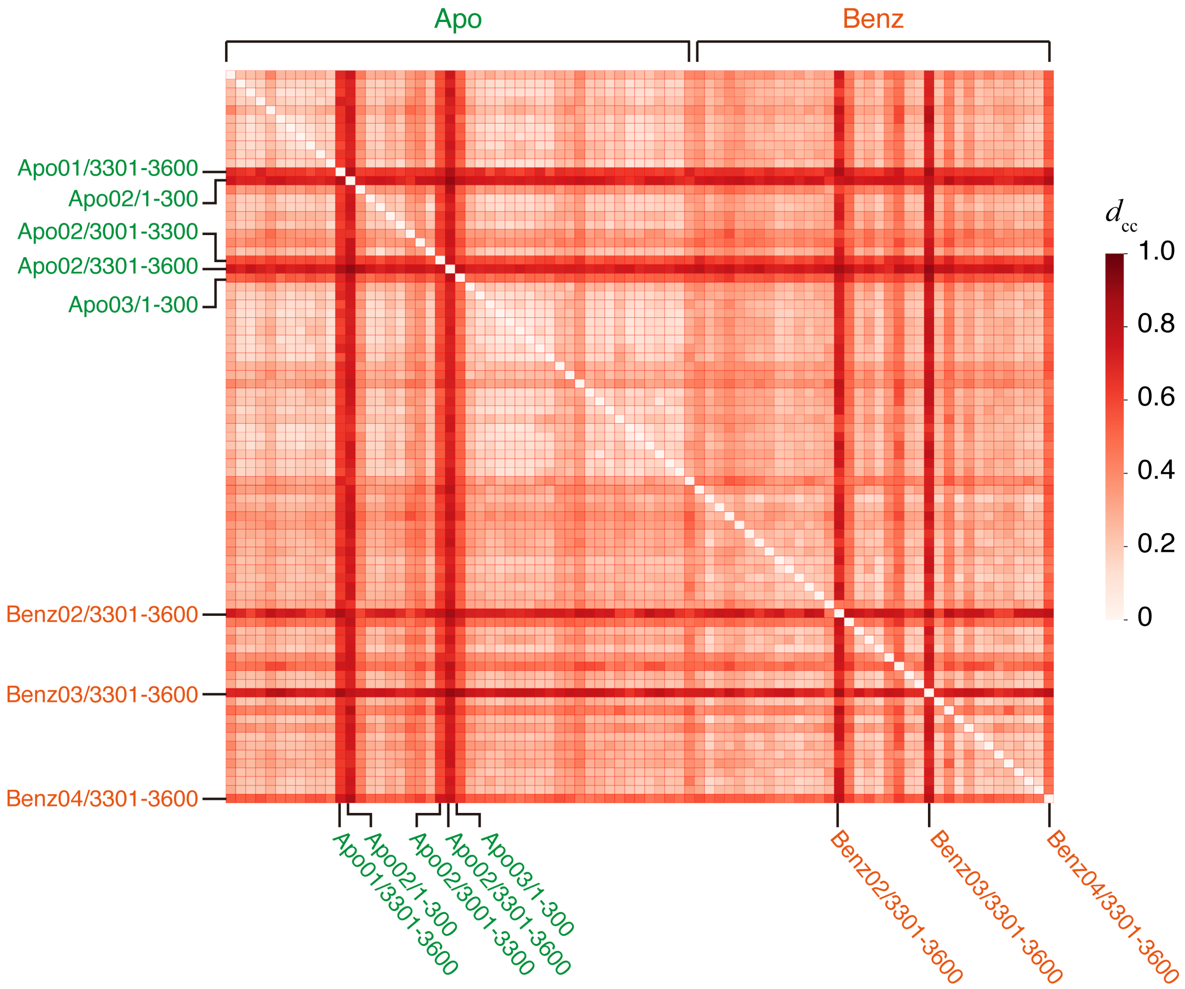


**Figure S12** Heat map diagram of $d_{CC}$ between all apo and benzamidine-bound trypsin chunks. In both vertical and horizontal axis, data label starts from the first 30° chunk of first apo-trypsin crystal and end with the last 30° chunk of last (fourth) benzamidine-bound trypsin crystal. The labelled chunk is not correlate with almost all of other chunks. The label indicates the data type, crystal ID, and the range of frame number.

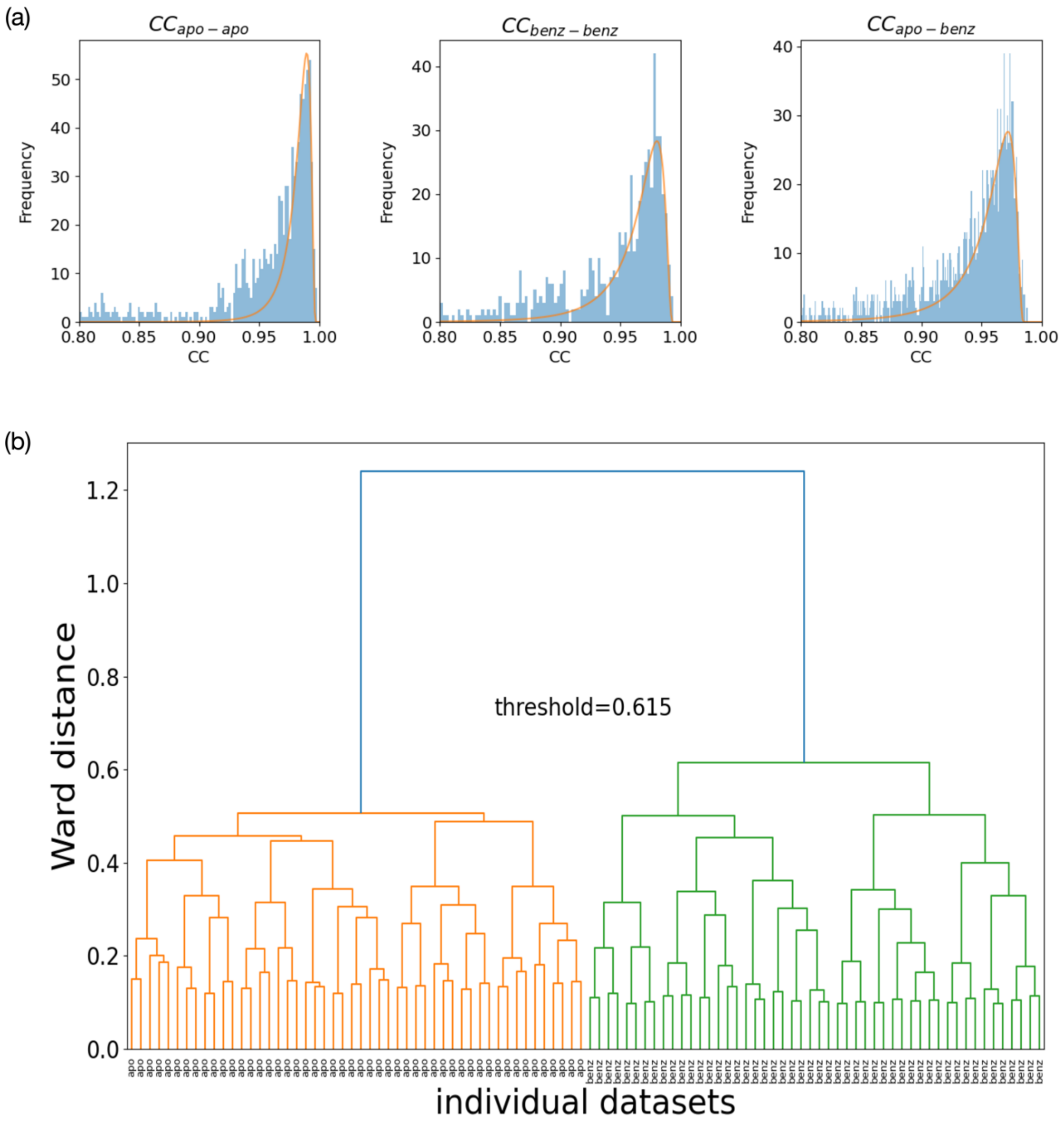


**Figure S13** HCA simulation based on the observed CC distributions using trypsin test case. (a)The log-normal distribution was fitted to each of observed $CC_{apo\text{-}apo}$, $CC_{benz\text{-}benz}$, $CC_{apo\text{-}benz}$. The fitted parameters (*s*, *loc*, *scale*) of log-normal distribution for $CC_{apo\text{-}apo}$, $CC_{benz\text{-}benz}$, and $CC_{apo\text{-}benz}$ were estimated as follows: (*s*, *loc*, *scale*) = (0.704, 0.003, 0.013), (0.765, 0.006, 0.025) and (0.787, 0.015, 0.025), respectively. Each model seems to fit the observed CC distribution well. (b) One of the dendrograms from the HCA simulation assuming a log−normal distribution of CC distribution, with 50 datasets each for the two trypsin structural polymorphs.

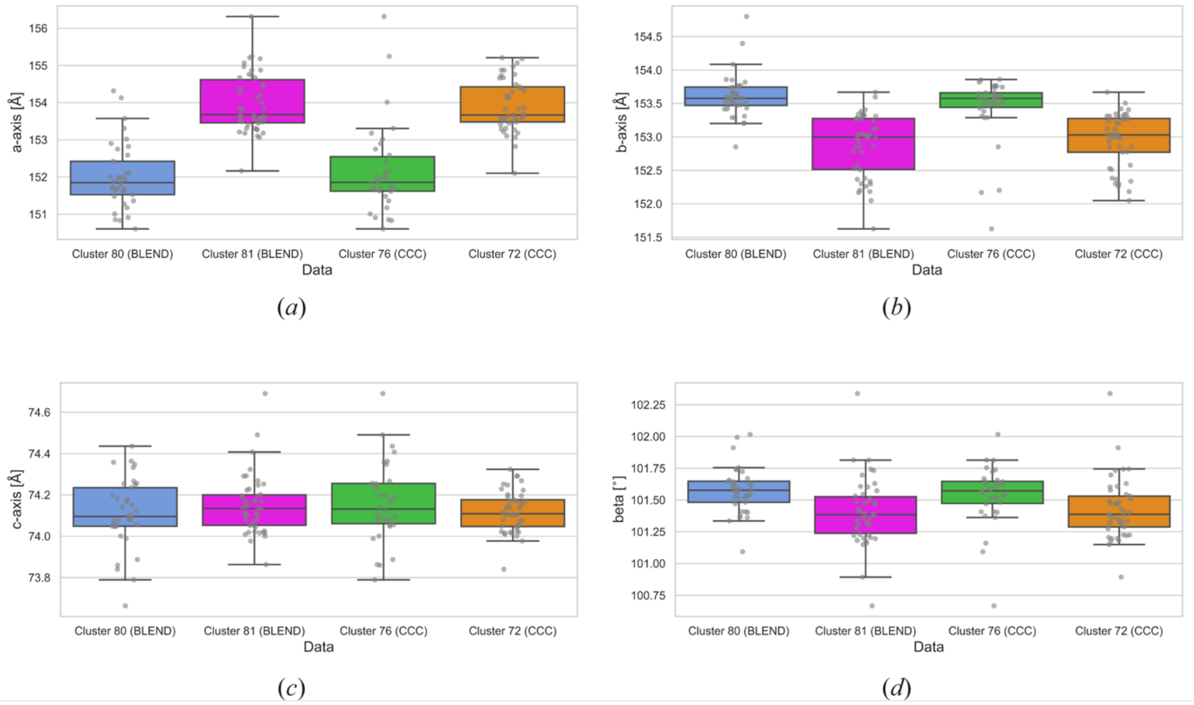


**Figure S14** Unit cell constants for the 30° chunks of Trn1-peptide complex. (a) a-axis, (b) b-axis, (c) c-axis, (d) beta angle. Distribution of the unit cell constants in clusters obtained by unit cell-based and intensity-based HCA were compared.

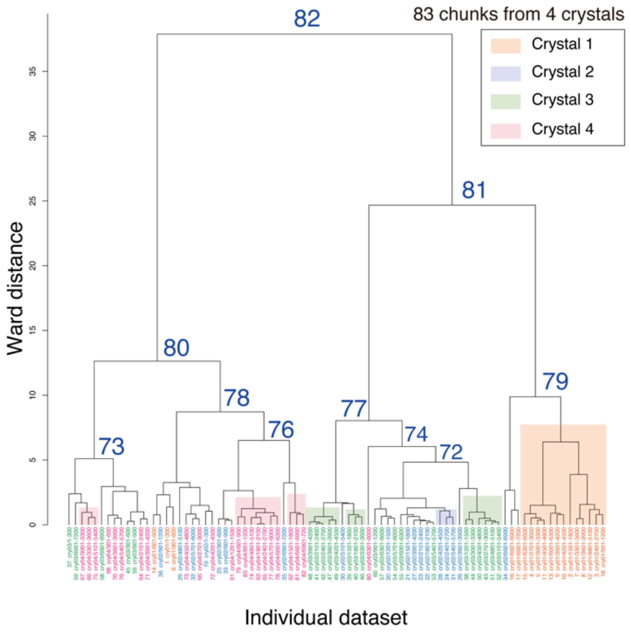


**Figure S15** Dendrogram from unit cell-based HCA on Trn1-NLS peptide complex. The data label are colored by the crystal ID; crystal 1: orange, crystal 2: blue, crystal 3: green, and crystal 4:

magenta. Cluster in the dendrogram is colored by the same color for the leaf label, when more than three chunks from the same crystal form cluster.

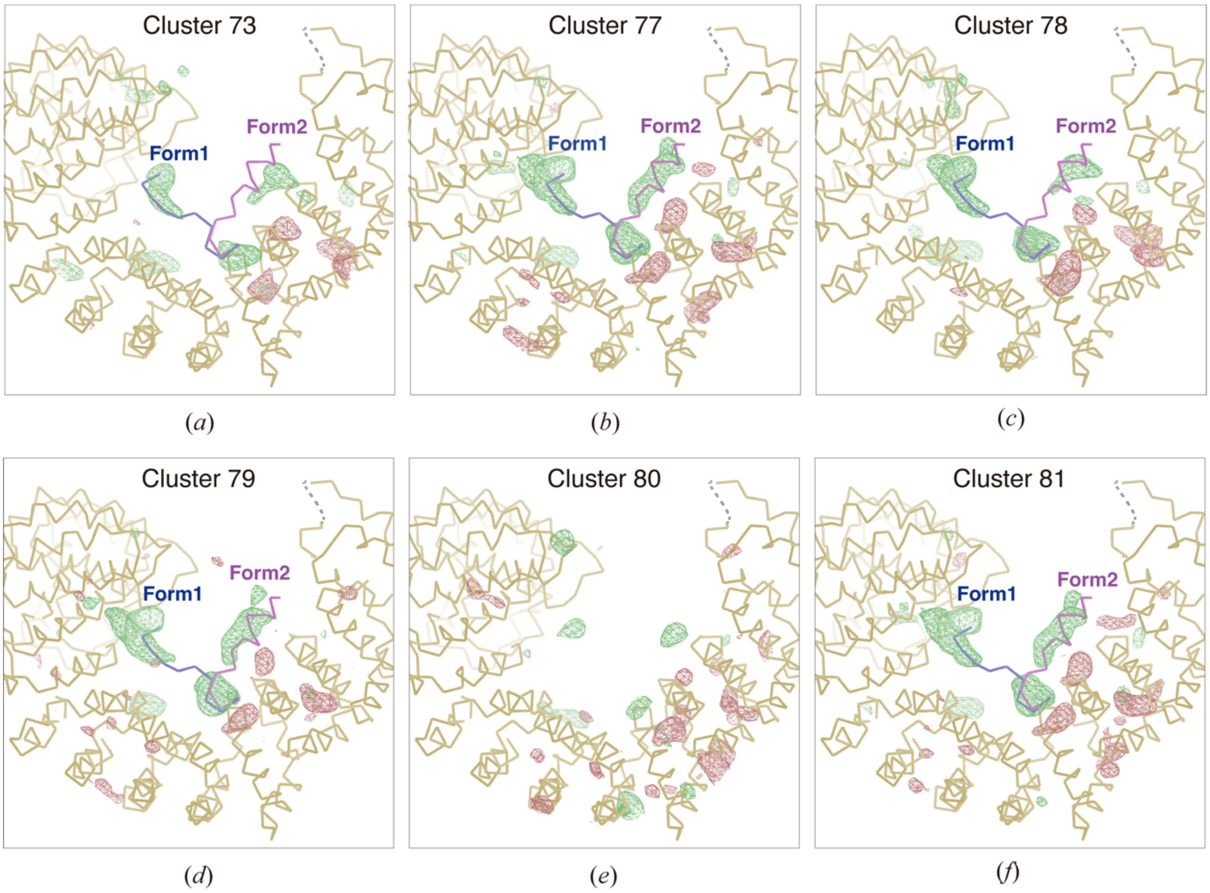


**Figure S16** Peptide-model omitted $F_o$-$F_c$ maps obtained at different clusters from the unit cell-based HCA on Trn1-peptide complex. Contour level for the $F_o$-$F_c$ map (green mesh: positive, and red mesh: negative) is set to 3.0σ. The maps were calculated without a model of binding peptide. Only the main chain for Trn1 and binding peptide is depicted. In contrast to the results from the intensity-based HCA, both peptide binding form were observed in a similar density. Figures were generated by *Coot*.

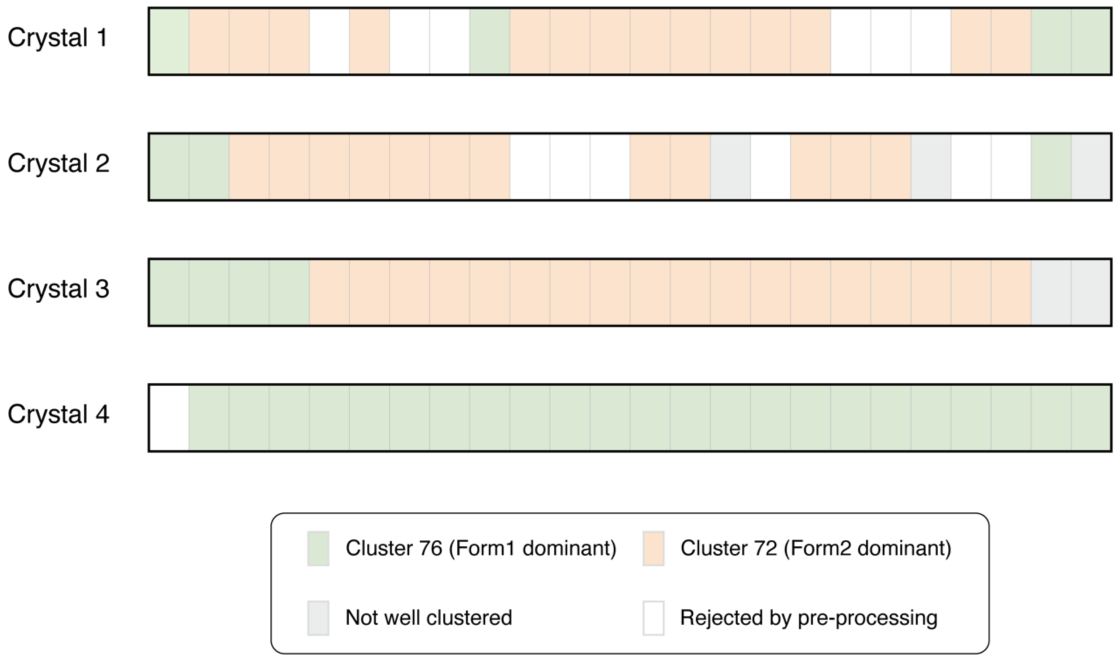


**Figure S17** Intra-crystal polymorphism implied by intensity-based HCA on Trn1-NLS peptide complex. Each 30° chunk was colored according to the result of intensity-based HCA; green: involved in Cluster 76 (Form1 dominant cluster), orange: involved in Cluster 72 (Form 2 dominant cluster), gray: not well clustered (outlier-like in the dendrogram), white: rejected in pre-processing by *KAMO* prior to the clustering.

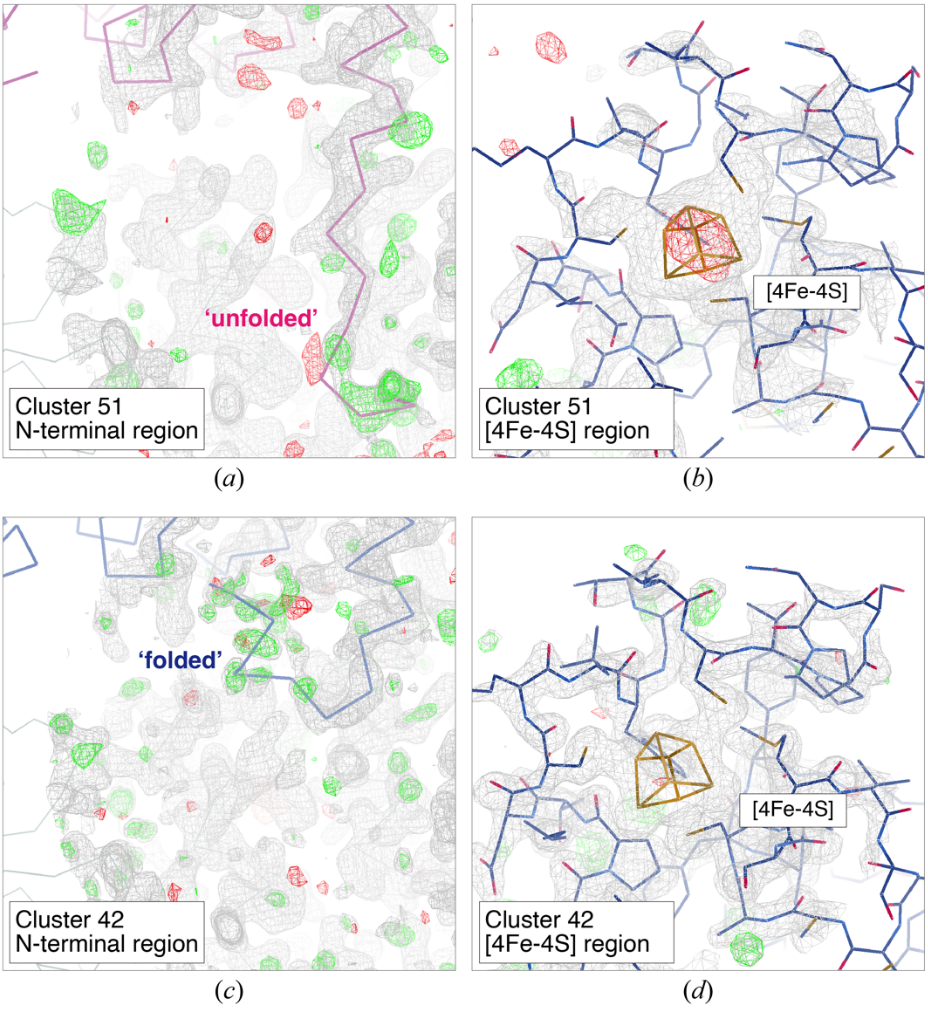


**Figure S18** Electron density maps around the N-terminal region and [4Fe-4S] cluster obtained from the merged data of *Aa*HypD-C360S at (a)-(b) Cluster 51 and (c)-(d) Cluster 42. Contour level for $2F_o$-$F_c$ map (blue mesh) is 1.0σ, except for [4Fe-4S] region in Cluster 42, where it is set to 1.5σ. The contour level for $F_o$-$F_c$ map (green mesh: positive, and red mesh: negative) is 3.0σ. Only the main chain of unfolded (purple) and folded models (pink) for the N-terminal region is depicted. The variable N-terminal region (Ser7-Tyr12) was omitted and the occupancy of [4Fe-4S] was set to 1.0 in map calculation. Figures were generated by *Coot*.

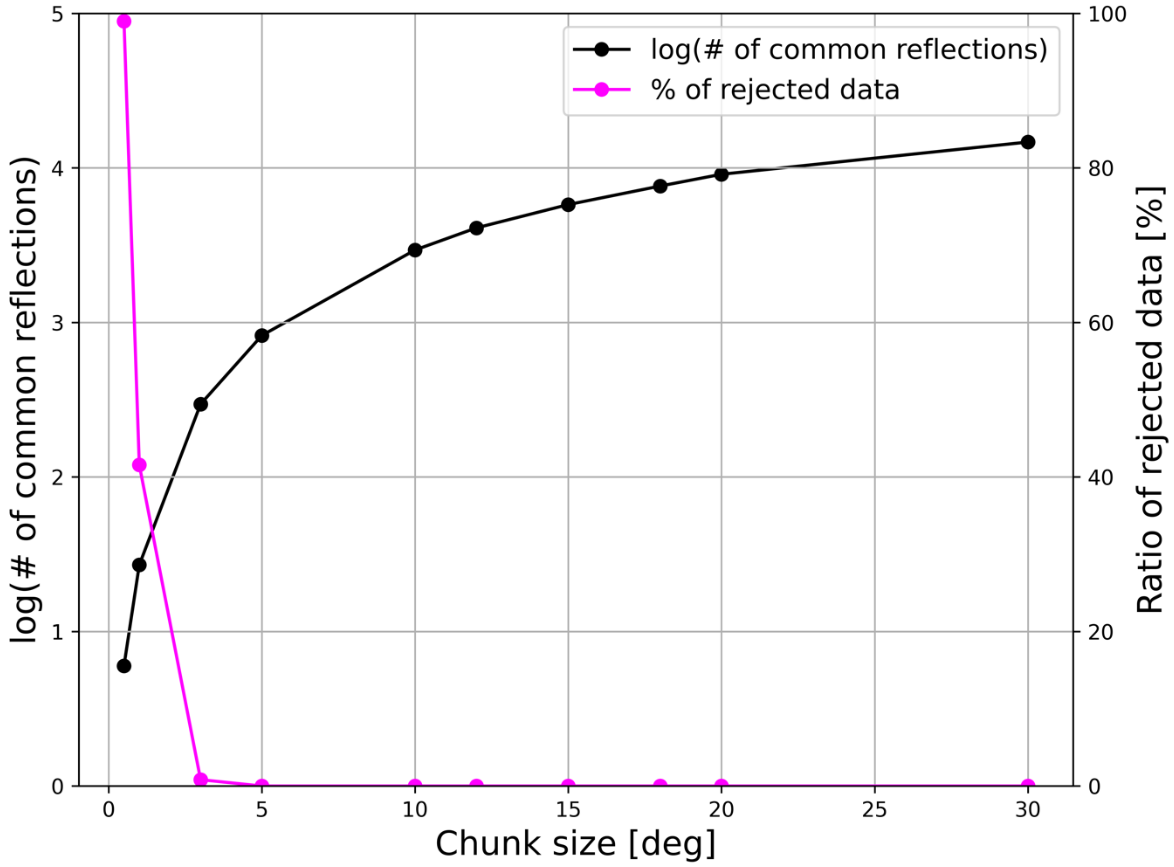


**Figure S19** The number of common reflections (black) and the ratio of rejected data (magenta) in the case of apo-trypsin. The higher resolution limit $d_{min}$ in CC calculation was set to 1.5 Å. Rejected data drastically increased when chunk size was set to below 3.0° due to the lack of common reflections to calculate CC value.

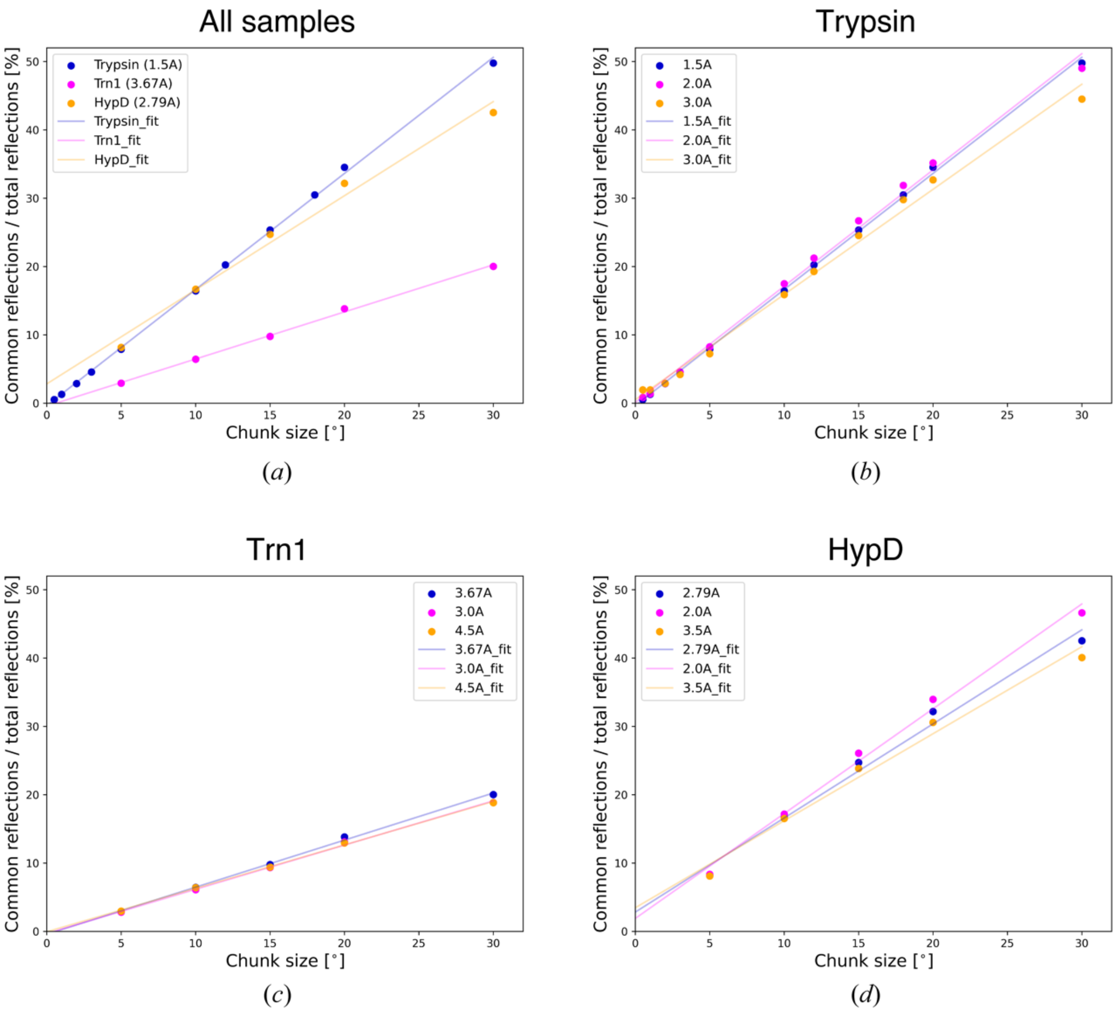


**Figure S20** Chunk size-dependent change of the fraction of common reflections in total reflections. (a) all samples (CC was calculated at the same cut-off resolution used in this study), (b)-(d) CC was calculated with different cut-off resolution. (b) trypsin with 1.5, 2.0, and 3.0 Å, (c) Trn1 with 3.0, 3.67, 4.5 Å, (d) HypD with 2.0, 2.79, 3.5 Å. The molecular weight of trypsin, Trn1, and *Aa*HypD-C360S are approximately 24 kDa, 98 kDa, and 43 kDa, respectively.

**Table S1** Summary of pre-processing prior to HCA by *KAMO*

| | | # of rejected chunks prior to HCA | | |
|---|---|---|---|---|
| Dataset | # of input chunks | Equivalent cell selection | Unit cell-based filtering | # of chunks used in HCA |
| Trypsin (apo + benz) | 96 | 2 | 11 | 83 |
| Trypsin (benz + tryp) | 96 | 2 | 10 | 84 |
| Trn1 | 96 | 7 | 6 | 83 |
| HypD | 72 | 2 | 3 | 67 |

**Table S2** Number of chunks in merged data at each node from unit cell-based clustering (apo-trypsin + benzamidine-bound trypsin)

| | Before outlier rejection | | After outlier rejection | | |
|---|---|---|---|---|---|
| Cluster # | Apo-trypsin | Benzamidine-bound | Apo-trypsin | Benzamidine-bound | Fo-Fc density |
| 67 | 10 | 2 | 5 (83.3%) | 1 (16.7%) | A |
| 72 | 11 | 0 | 8 (100%) | 0 (0%) | A |
| 74 | 17 | 2 | 8 (88.9%) | 1 (11.1%) | A |
| 76 | 14 | 0 | 9 (100%) | 0 (0%) | A |
| 77 | 3 | 18 | 2 (15.4%) | 11 (84.6%) | B |
| 78 | 2 | 16 | 0 (0%) | 10 (100%) | B |
| 79 | 25 | 0 | 20 (100%) | 0 (0%) | A |
| 80 | 20 | 20 | 14 (46.7%) | 16 (53.3%) | B? |

| | | | | | |
|---|---|---|---|---|---|
| 81 | 45 | 20 | 33 (71.7%) | 13 (28.3%) | A |
| 82 | 47 | 36 | 35 (59.3%) | 24 (40.7%) | A |

* A: apo-trypsin, B: benzamidine-bound trypsin (? indicates uncertain assignment)

**Table S3** Number of chunks in merged data at each node from unit cell-based clustering (benzamidine-bound trypsin + tryptamine-bound trypsin)

| | Before outlier rejection | | After outlier rejection | | |
|---|---|---|---|---|---|
| Cluster # | Benzamidine-bound | Tryptamine-bound | Benzamidine-bound | Tryptamine-bound | Fo-Fc density |
| 70 | 0 | 14 | 0 (0%) | 8 (100%) | T |
| 74 | 11 | 0 | 8 (100%) | 0 (0%) | B |
| 76 | 0 | 21 | 0 (0%) | 17 (100%) | T |
| 77 | 13 | 7 | 9 (69.2%) | 4 (30.8%) | B |
| 78 | 11 | 3 | 5 (71.4%) | 2 (28.6%) | B |
| 79 | 1 | 29 | 0 (0%) | 21 (100%) | T |
| 80 | 13 | 16 | 6 (35.0%) | 14 (65.0%) | indistinguishable |
| 81 | 24 | 16 | 18 (58.1%) | 13 (41.9%) | B? |
| 82 | 12 | 32 | 5 (18.5%) | 22 (81.5%) | T |
| 83 | 36 | 48 | 23 (37.7%) | 38 (62.3%) | T |

* B: benzamidine-bound trypsin, T: tryptamine-bound trypsin (? indicates uncertain assignment)

**Table S4** Data statistics for merged data obtained from intensity-based clustering (apo + benzamidine-bound trypsin)

| Cluster # | # of chunks | Resolution [Å] | Redundancy | Completeness [%] | $R_{meas}$ [%] | <I/σI> | $CC_{1/2}$ |
|---|---|---|---|---|---|---|---|
| 63 | 27 | 1.20 | 24.2 (26.9/11.16) | 99.2 (99.7/95.2) | 4.7 (3.5/48.1) | 39.36 (86.58/5.39) | 99.9 (99.8/93.6) |
| 65 | 29 | 1.20 | 26.0 (29.0/11.97) | 99.2 (99.7/95.3) | 5.0 (3.6/53.2) | 39.59 (87.31/5.34) | 99.9 (99.8/93.2) |
| 67 | 18 | 1.20 | 16.8 (19.5/7.90) | 98.8 (99.9/92.8) | 7.6 (6.5/97.9) | 22.29 (49.38/2.36) | 99.9 (99.7/73.6) |
| 69 | 28 | 1.20 | 25.5 (28.3/11.74) | 99.2 (99.8/95.2) | 4.9 (3.7/51.9) | 39.6 (86.76/5.46) | 99.9 (99.8/93.7) |
| 70 | 23 | 1.21 | 21.6 (24.8/10.85) | 99.2 (99.9/95.2) | 12.2 (11.5/99.6) | 23.21 (50.56/2.83) | 99.8 (99.5/78.5) |
| 71 | 32 | 1.20 | 28.7 (32.2/13.19) | 99.9 (99.8/92.3) | 5.7 (4.5/57.8) | 39.2 (87.7/5.37) | 99.9 (99.8/92.3) |
| 72 | 18 | 1.20 | 16.9 (19.6/7.90) | 98.9 (99.9/93.4) | 7.5 (6.6/93.5) | 22.57 (48.39/2.71) | 99.8 (99.5/73.6) |
| 76 | 31 | 1.20 | 28.2 (31.5/12.98) | 99.2 (99.7/95.2) | 5.2 (4.1/54.2) | 41.38 (86.30/6.6) | 99.9 (99.8/92.9) |
| 81 | 53 | 1.20 | 48.7 (55.3/21.76) | 99.9 (99.8/92.5) | 11.9 (10.9/79.3) | 31.69 (59.1/5.94) | 99.9 (99.8/92.5) |
| 82 | 59 | 1.20 | 54.6 (62.3/24.38) | 99.8 (99.9/98.9) | 15.3 (14.3/84.3) | 31.79 (58.8/6.13) | 99.8 (99.6/92.3) |

* Values in parentheses indicate inner/outer resolution shell. Number of chunks is the value based on the final merged datasets after outlier rejection process by *KAMO*.

**Table S5** Data statistics for merged data obtained from intensity-based clustering (benzamidine-bound + tryptamine-bound trypsin)

| Cluster # | # of chunks | Resolution [Å] | Redundancy | Completeness [%] | $R_{meas}$ [%] | <I/σI> | $CC_{1/2}$ |
|---|---|---|---|---|---|---|---|
| 58 | 13 | 1.20 | 12.9 (16.6/6.06) | 94.8 (86.8/88.8) | 5.2 (5.4/37.3) | 27.18 (54.37/5.35) | 99.9 (99.7/93.2) |
| 62 | 12 | 1.22 | 11.7 (13.2/6.37) | 99.1 (99.8/94.7) | 11.4 (9.9/117.2) | 17.07 (45.64/1.71) | 99.8 (99.6/63.7) |
| 66 | 18 | 1.21 | 17.3 (19.9/8.75) | 99.2 (99.9/95.1) | 9.6 (8.8/91.7) | 22.69 (49.38/2.68) | 99.9 (99.6/75.1) |
| 67 | 24 | 1.20 | 22.9 (26.6/10.61) | 99.0 (99.9/93.9) | 5.3 (5.4/37.4) | 39.78 (77.74/7.15) | 99.9 (99.8/95.5) |
| 72 | 26 | 1.20 | 25.0 (29.1/11.58) | 99.0 (99.9/94.0) | 5.5 (5.6/37.1) | 40.80 (78.46/7.64) | 99.9 (99.9/95.8) |
| 74 | 18 | 1.21 | 17.4 (19.9/8.75) | 99.2 (99.9/95.1) | 9.6 (8.8/91.7) | 22.69 (49.36/2.68) | 99.9 (99.7/74.5) |
| 75 | 21 | 1.21 | 22.5 (25.9/11.23) | 99.3 (99.9/95.9) | 12.9 (12.4/106.2) | 23.04 (51.08/2.79) | 99.9 (99.6/74.5) |
| 79 | 17 | 1.21 | 25.4 (29.1/12.76) | 99.0 (99.9/94.0) | 22.43 (21.1/112.7) | 22.12 (45.86/3.37) | 99.7 (99.4/69.1) |
| 82 | 15 | 1.21 | 20.0 (23.0/10.09) | 99.2 (99.9/95.3) | 8.0 (6.9/105.4) | 23.66 (53.61/2.58) | 99.8 (99.6/69.7) |
| 83 | 41 | 1.20 | 41.3 (47.7/19.11) | 99.2 (99.9/95.1) | 10.9 (8.9/68.1) | 27.22 (50.26/6.4) | 99.9 (99.8/91.9) |

* Values in parentheses indicate inner/outer resolution shell. Number of chunks is the value based on the final merged datasets after outlier rejection process by *KAMO*.

**Table S6** Data statistics for merged data obtained from intensity-based clustering (Trn1-NLS peptide complex)

| Cluster # | # of chunks | Resolution [Å] | Redundancy | Completeness [%] | $R_{meas}$ [%] | <I/σI> | $CC_{1/2}$ |
|---|---|---|---|---|---|---|---|
| 72 | 44 | 3.89 | 21.6 (22.5/22.22) | 99.9 (98.4/100) | 10.1 (5.1/319.9) | 17.79 (67.16/1.66) | 100 (99.9/78.1) |
| 76 | 34 | 4.43 | 15.8 (17.1/16.09) | 99.8 (97.6/99.6) | 11.1 (4.8/327.2) | 15.77 (71.34/1.1) | 100 (100/58.5) |
| 78 | 35 | 4.43 | 16.3 (17.6/16.57) | 99.8 (97.6/99.7) | 11.3 (4.9/337.0) | 15.83 (71.47/1.1) | 100 (100/57.9) |
| 81 | 79 | 3.88 | 39.1 (40.4/39.97) | 99.9 (98.2/101) | 19.5 (7.2/1551.1) | 19.2 (75.02/1.72) | 100 (100/77.1) |
| 82 | 83 | 3.88 | 41.2 (42.5/42.01) | 99.9 (98.2/100) | 20.6 (7.5/2945.6) | 19.33 (76.11/1.72) | 100 (100/79.7) |

* Values in parentheses indicate inner/outer resolution shell. Number of chunks is the value based on the final merged datasets after outlier rejection process by *KAMO*.

**Table S7** Data statistics for merged data obtained from intensity-based clustering (*Aa*HypD-C360S).

| Cluster # | # of chunks | Resolution [Å] | Redundancy | Completeness [%] | $R_{meas}$ [%] | <I/σI> | $CC_{1/2}$ |
|---|---|---|---|---|---|---|---|
| 42 | 12 | 1.85 | 9.1 (8.3/9.4) | 100 (99.6/100) | 10.7 (8.7/27.3) | 17.98 (31.73/7.17) | 99.8 (99.7/98.8) |
| 47 | 2 | 1.85 | 5.6 (3.0/2.6) | 79.7 (64.1/84.1) | 10.2 (5.6/102.0) | 4.4 (9.18/1.11) | 99.6 (99.6/63.1) |
| 50 | 10 | 1.88 | 7.5 (7.0/7.5) | 99.9 (99.3/99.9) | 13.4 (4.2/316.7) | 13.17 (45.13/1.15) | 99.9 (99.9/59.0) |
| 51 | 13 | 2.02 | 9.5 (8.9/9.6) | 99.9 (99.2/100) | 20.7 (5.2/397.4) | 11.17 (43.2/1.08) | 99.8 (99.9/55.0) |
| 52 | 6 | 2.20 | 5.9 (5.7/5.7) | 99.8 (98.1/99.9) | 24.2 (8.3/130.4) | 5.58 (17.03/1.25) | 99.0 (99.5/54.6) |
| 54 | 5 | 1.94 | 4.8 (4.9/4.7) | 98.9 (95.0/99.4) | 26.0 (12.7/125.6) | 3.42 (10.22/0.70) | 98.2 (98.3/57.6) |
| 61 | 23 | 1.85 | 16.4 (15.2/16.4) | 100 (99.5/100) | 32.1 (7.6/N.D.) | 9.97 (36.67/0) | 99.8 (99.9/63.8) |
| 63 | 17 | 1.96 | 14.6 (14.4/14.1) | 100 (99.5/100) | 52.9 (16.7/233.6) | 4.09 (14.02/0.74) | 97.3 (99.2/52.5) |
| 64 | 27 | 1.85 | 8.2 (7.6/8.3) | 100 (99.6/100) | 13.7 (6.6/112.2) | 18.05 (32.59/6.83) | 99.5 (98.7/97.8) |
| 65 | 44 | 1.85 | 28.1 | 100 | 47.2 | 17.03 | 98.7 |

| | | | | | | | |
|---|---|---|---|---|---|---|---|
| | | | (26.8/27.7) | (99.6/100) | (30.6/167.1) | (31.47/6.95) | (96.7/96.3) |
| 66 | 67 | 1.85 | 49.8 | 100 | 86.8 | 7.17 | 98.7 |
| | | | (47.0/48.5) | (99.6/100) | (36.2/3883.7) | (18.06/3.1) | (99.3/91.2) |

* Values in inner/outer resolution shell are indicted in parentheses. N.D. = Not determined by XSCALE. Number of chunks is the value based on the final merged datasets after outlier rejection process by *KAMO*.
